# Hierarchical Immune Suppressive Functions of Regulatory T Cells Built on Mechanical Force

**DOI:** 10.64898/2026.05.18.725872

**Authors:** Xinyi Liu, Cheng Huang, Xiaobo Wang, Jie Kang, Chunlin Zou, Yanni Xu, Longyan Wu, Yan Shi

## Abstract

Regulatory T (Treg) cells maintain immune tolerance via contact-dependent inhibition and soluble mediators. While canonical suppressive mechanisms are well characterized, their hierarchical spatiotemporal organization remains unclear. We previously showed Foxp3 represses ER Ca^2+^ channel RyR2, reducing cytoplasmic Ca^2+^ to inactivate m-Calpain (Calpain-2) and stabilize high-affinity LFA-1-mediated Treg–dendritic cell (DC) adhesion—an early step blocking DC antigen presentation. Using m-Calpain as a molecular switch, we generated mechanically deficient Tregs (TregFD) with constitutively active m-Calpain that abrogated high-force Treg-DC adhesion. TregFD retained a near-wild-type transcriptome (differential expression limited to adhesion pathways), yet failed to suppress autoimmunity or DC-driven T cell proliferation. Stable adhesion was required for localized TGF-β/IL-10 delivery to DCs. Single-cell RNA sequencing of rescued Scurfy mouse lymphoid tissues revealed organ-specific division of labor: non-mechanical modules support homeostasis/tissue repair, while mechanical modules dominate epithelial barrier/antigen presentation suppression. Combined TregFD and RyR2-deficient Tconv transfer restored wild-type Treg activity, demonstrating module synergism. These findings establish a mechanical force-centered, two-tiered hierarchical model of Treg suppression, providing a framework for targeting Treg mechanics in autoimmunity and inflammation.

## Main Text

Regulatory T (Treg) cells are the central arbiters of immune tolerance and tissue homeostasis (*1, 2*). Since the discovery of Foxp3 as the lineage-defining transcription factor (*3-5*), intensive investigation has defined an array of effector mechanisms that contribute to Treg suppression, including inhibitory cytokine secretion (TGF-β, IL-10, IL-35), metabolic disruption (adenosine via CD39/CD73), cytokine absorption (IL-2 deprivation), cytolytic activity (granzyme/perforin), and surface receptor–mediated inhibition (CTLA-4, PD-1, LAG-3, TIGIT) (*6-11*). These suppressive pathways are not unique to Tregs or controlled by Foxp3, yet only Tregs exhibit dominant suppressive activity in vivo. This points to a conceptual deficit: In contrast to conventional T cells whereby all those effector mechanisms are also available, how does Foxp3 organize these diverse modalities to produce robust, context-appropriate, and Treg-specific suppression?

Here we provide comprehensive functional, molecular, imaging, transcriptomic, and in vivo evidence that Treg mechanical suppression orchestrated by Foxp3 is the structural basis for other immune suppressive properties in addition to its blockage of DC antigen presentation. We demonstrate that m-Calpain overexpression selectively abolishes stable Treg–DC adhesion while preserving Treg development, homeostasis, phenotype, trafficking, and soluble inhibitory functions. Using multiple autoimmune disease models, we show that mechanical adhesion is required for full Treg suppressive activity in vivo. Mechanistically, mechanical adhesion licenses rapid, localized inhibitory signaling via TGF-β and IL-10 to DCs, revealing a hierarchical organization where mechanics precede and enable soluble suppression. Transcriptomic and single-cell analyses further reveal a tissue-specific division of labor between mechanical and non-mechanical suppressive arms. Together, these findings redefine the core logic of Treg suppression, establish mechanical regulation as a foundational pillar of Treg biology, and nominate m-Calpain as a promising therapeutic target for modulating Treg activity in autoimmunity.

### Engineering and Validation of Mechanically Deficient Tregs (TregFD)

In vivo imaging shows Tregs form stable dendritic cell (DC) interactions to suppress antigen presentation and effector T cell priming (*12-15*). Our prior work identified a critical mechanical component: Tregs exert adhesive forces via LFA-1–ICAM-1 to inactivate DCs (*16*). The LFA-1-mediated binding is pivotal for Treg function has been verified in in vitro suppression assay (fig. S1A). We further delineated a Foxp3–RyR2–Ca^2^□–m-Calpain– Talin-1–LFA-1 signaling cascade that underpins this mechanical phenotype (*17*). Foxp3 represses ER Ca^2^□ channel RyR2, reducing Ca^2^□ oscillations and m-Calpain activation; inactive m-Calpain preserves Talin-1, stabilizing LFA-1 high affinity DC binding. Despite this, causal links between Treg mechanical function and in vivo suppression remain elusive. To address this, we engineered Treg Force Deficient (TregFD) cells and mice via Treg-specific overexpression of a Ca^2^□-insensitive, constitutively active m-Calpain mutant— uncoupling mechanical adhesion from other suppressive modules to define its role in Treg function. CA-m-Calpain was constructed by mutating Ser369 and Thr370 to alanine (ST369/370AA), rendering the protease independent of Ca^2^□ regulation (*18*). We expressed this construct in primary mouse Tregs via lentiviral transduction, achieving approximately 32-fold overexpression of m-Calpain mRNA relative to control Tregs (Fig. 1A). To quantify adhesive force at the single-cell level, we used atomic force microscopy–based single-cell force spectroscopy (AFM-SCFS) (*19, 20*). Wild-type (WT) Tregs exhibited strong binding to DC2.4 DCs, with a mean adhesion force of 1.1 ± 0.1 nN. By contrast, TregFD cells showed a mean adhesion force of 0.75 ± 0.05 nN, nearly identical to that of Tconv cells (0.7 ± 0.1 nN; Fig. 1B). Overexpression of μ-Calpain (Calpain-1) did not alter adhesion force in line with our previous work (*16*), indicating subtype specificity for m-Calpain in this mechanical pathway (fig. S1B).

**Fig. 1.**
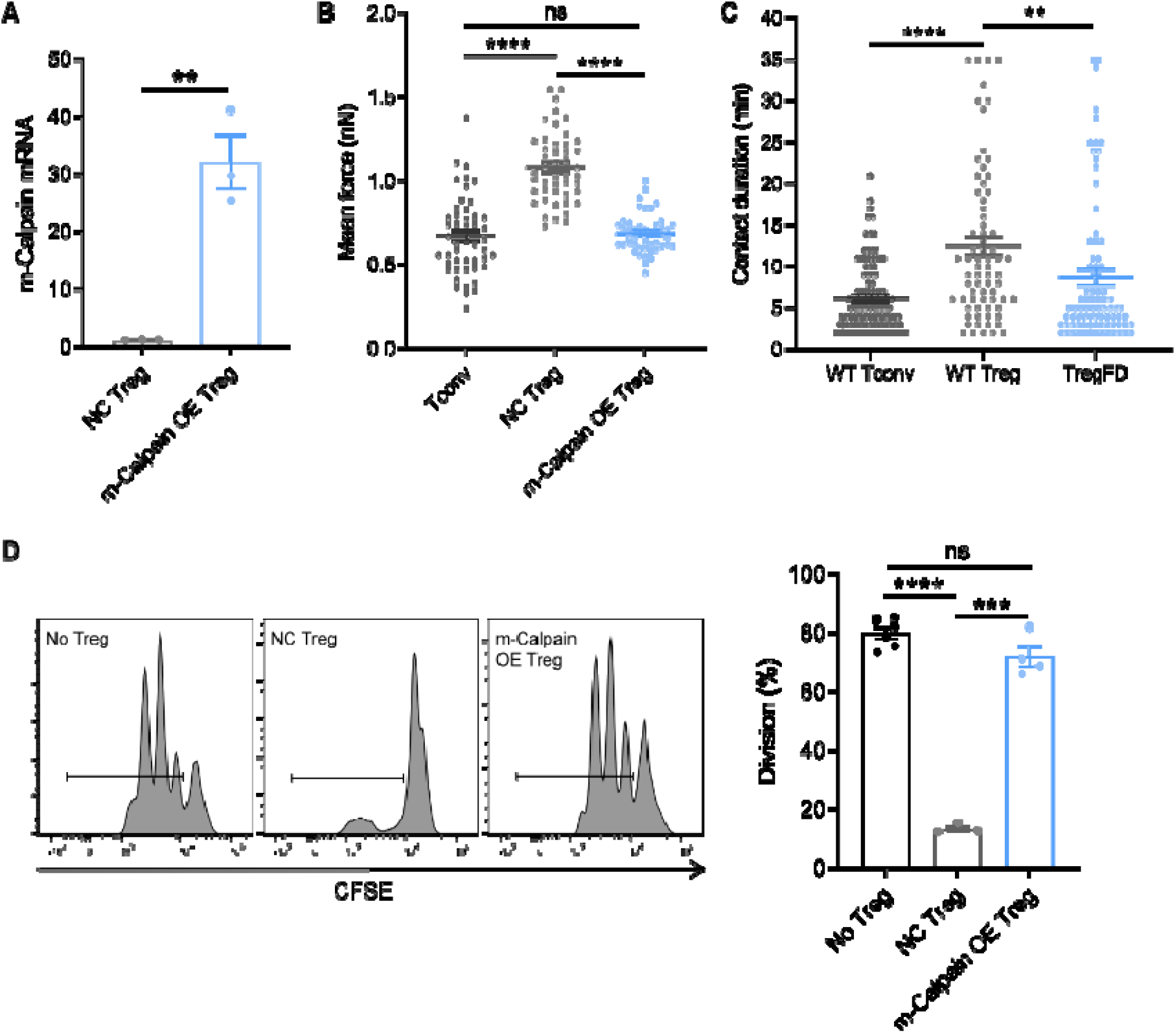
m-Calpain Overexpression Abolishes Treg–DC Mechanical Adhesion and Suppressive Function. (**A**) Relative mRNA expression of Capn2 (m-Calpain) in control Tregs and Tregs overexpressing constitutively active m-Calpain (TregFD). Data are mean ± SEM; n = 3 biological replicates; **p < 0.01, two-tailed Student’s t-test. (**B**) Single-cell adhesion forces between DC2.4 DCs and Tconv, WT Treg, or TregFD cells measured by AFM-SCFS. Each dot represents one measurement; n ≥ 15 per group; mean ± SEM; ****p < 0.0001, one-way ANOVA with Tukey’s multiple comparisons test. (**C**) In vivo contact duration between T cells and CD11c□DCs in lymph nodes measured by intravital confocal microscopy. n = 5 mice per group; mean ± SEM; **p < 0.01, ****p < 0.0001, one-way ANOVA with Tukey’s multiple comparisons test. (**D**) In vitro suppression of OT-II CD4□T cell proliferation by WT Treg or TregFD. Representative CFSE dilution plots (left) and quantification (right). n = 3 independent experiments; mean ± SEM; ***p < 0.001, ****p < 0.0001, one-way ANOVA with Tukey’s multiple comparisons test.

We next performed intravital confocal imaging of inguinal lymph nodes to measure Treg–DC contact dynamics in vivo per our published protocol (*17*). DCs were labeled via CD11c-GFP, and T cells were labeled with CellTrace Far Red. WT Tregs formed stable contacts with DCs lasting 12.3 ± 1.5 minutes. TregFD–DC contact duration was reduced to 8.8 ± 1.1 minutes, comparable to Tconv–DC contacts (6.1 ± 0.9 min; Fig. 1C). These data confirm that m-Calpain overexpression abolishes both the high adhesive force and prolonged contact duration that define Treg–DC mechanical interactions. To assess functional suppression, we performed in vitro CFSE-based proliferation assays using OT-II CD4□ T cells as responders. WT Tregs strongly inhibited OT-II proliferation, while TregFD cells lost nearly all suppressive capacity (Fig. 1D). Together, these results demonstrate that m-Calpain overexpression generates Tregs that are selectively deficient in mechanical adhesion and contact-dependent suppression.

### Generation and Characterization of TregFD Conditional Transgenic Mice

A Treg-specific conditional m-Calpain overexpression mouse line was produced using CRISPR/Cas9-mediated knock-in at the H11 safe-harbor locus. The construct contained a CAG promoter, a LoxP-Stop-LoxP (LSL) cassette, the CA-m-Calpain coding sequence, and a CFP fluorescent reporter (Fig. 2A). We crossed these mice to Foxp3-Cre-YFP mice to generate TregFD animals, in which CA-m-Calpain is expressed specifically in Foxp3□ Tregs. We also generated tamoxifen-inducible iTregFD mice by crossing to Foxp3-Cre-ERT2 to avoid potential developmental effects. Flow cytometry sorting of CFP□ TregFD cells confirmed ∼7-fold overexpression of m-Calpain mRNA (Fig. 2B) and strong transgene protein expression by Western blot (Fig. 2C and fig. S2A). In iTregFD mice, CFP expression increased gradually over 96 hours following 4-OHT treatment, indicating timed induction (fig. S2B). To confirm protease activity, we used a CMAC-calpain substrate cleavage assay. TregFD cells exhibited a marked rightward shift in fluorescence intensity relative to WT Tregs, indicating elevated calpain activity (Fig. 2D) (*21, 22*). This activity was insensitive to the RyR2 inhibitor JTV-519 (*23, 24*), confirming constitutive Ca^2^□-independent proteolysis (fig. S2C). TregFD animals were born at Mendelian ratios, exhibited normal growth, body weight, and lifespan, and showed no gross developmental abnormalities. Treg proliferation in response to anti-CD3/CD28 stimulation was comparable to WT Tregs (fig. S3A). TregFD retained high expression of IL-10 and TGF-β and low IL-2 secretion, consistent with WT Treg (fig. S3B). No differences in CD25, CD39, GITR, CTLA-4, or PD-1 between resting/activated TregFD and WT Tregs were found (fig. S3C). CD4□, CD8□, Foxp3□ Tregs, B cells, DCs, NK cells, and myeloid cells in the spleen, thymus, and lymph nodes were normal (fig. S3, D and G). In vivo trafficking assays using the Lumina III small-animal imaging system showed identical tissue distribution of TregFD and WT Tregs across lymphoid and non-lymphoid organs (fig. S3, E and F). Bone marrow chimera experiments confirmed that m-Calpain overexpression did not affect hematopoietic development (fig. S3H). TCR Vβ repertoire analysis revealed no alterations in diversity, indicating intact thymic selection (fig. S3I). Collectively, these data demonstrate that m-Calpain overexpression does not alter Treg development, homeostasis, phenotype, or trafficking—only mechanical adhesive function.

**Fig. 2.**
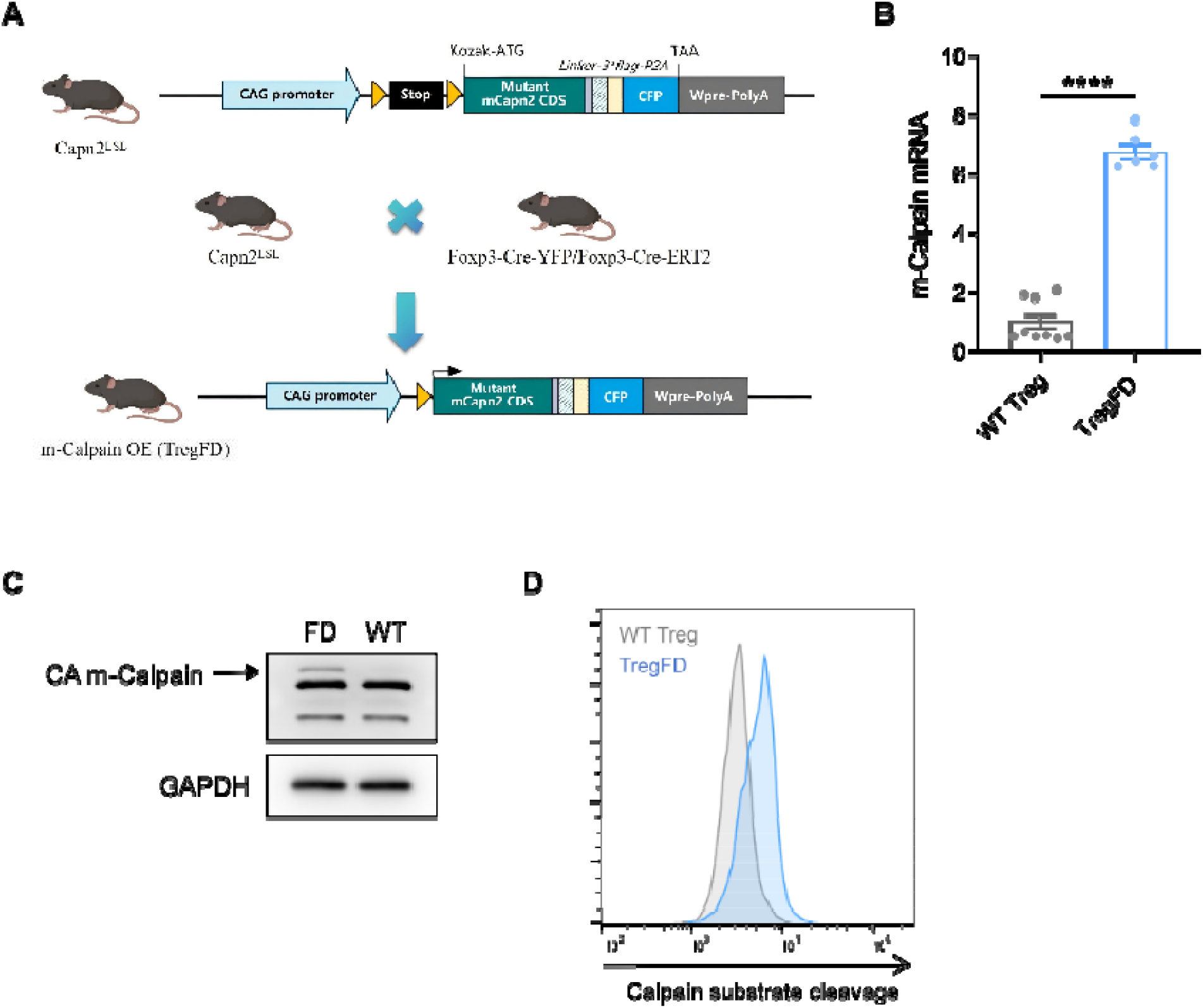
Generation and Validation of Treg-Specific m-Calpain Transgenic Mice (TregFD). (**A**) Schematic of the CAG-LSL-CA-m-Calpain-CFP construct knocked into the mouse H11 safe-harbor locus. (**B**) Relative Capn2 mRNA expression in sorted WT Treg and TregFD cells. N = 3 biological replicates; mean ± SEM; ****p < 0.0001, two-tailed Student’s t-test. (**C**) Western blot analysis of m-Calpain protein in WT Treg and TregFD cells. GAPDH served as loading control. Representative of three independent experiments. (**D**) Flow cytometry analysis of calpain substrate cleavage in WT Treg and TregFD cells. n = 3.

### TregFD Loses the Ability to Block DC-Mediated Antigen Presentation

With AFM/SCFS, the adhesive force of the two TregFD variants to DCs were more similar to those of Tconv cells, but significantly lower than those of WT Treg cells, indicating successful construction of TregFD (Fig. 3A). To confirm that TregFD cannot disrupt DC–Tconv interactions in vivo, we performed competitive intravital imaging assays. OT-II T cells were labeled and co-injected with unlabeled Tconv, WT Treg, TregFD, or RyR2 CKO Tconv cells into CD11c-GFP mice. WT Tregs strongly reduced DC–OT-II contact duration, while TregFD and Tconv had no effect (Fig. 3B). In vitro suppression assays recapitulated this finding: TregFD failed to inhibit OT-II proliferation, confirming loss of function (Fig. 3C). To define the transcriptomic feature alteration of m-Calpain overexpression, we performed bulk RNA sequencing (RNA-seq) on resting and activated Tconv, WT Treg, and TregFD cells. Principal component analysis (PCA) showed that TregFD clustered tightly with WT Tregs and distinctly from Tconvs, confirming preserved Treg transcriptional identity (Fig. 3D). Differential expression analysis identified only a small number of differentially expressed genes (DEGs) between activated WT Treg and TregFD, with *Capn2* being the most significantly altered (fig. S4A). Gene Ontology (GO) and Kyoto Encyclopedia of Genes and Genomes (KEGG) pathway enrichment revealed that DEGs were strongly enriched for cell adhesion, cytoskeletal organization, focal adhesion, and membrane component pathways (Fig. 3E). Heatmap analysis showed no significant differences in canonical Treg suppressor genes, including *Ctla4, Pdcd1, Il2ra, Tgfb1, Ebi3*, and *Ikzf2* (Fig. 3F). Gene Set Enrichment Analysis (GSEA) showed upregulation of immune synapse and cell adhesion gene sets in TregFD (fig. S4B), consistent with a compensatory response to reduced mechanical adhesion. Under the same conditions (resting or activated), the levels of FOX transcription factors in Treg and TregFD cells were similar, but they were far different from those in Tconv cells (fig. S4C). These transcriptomic data confirm that TregFD is a pure mechanical-deficient model, with no global perturbation of Treg identity or function.

**Fig. 3.**
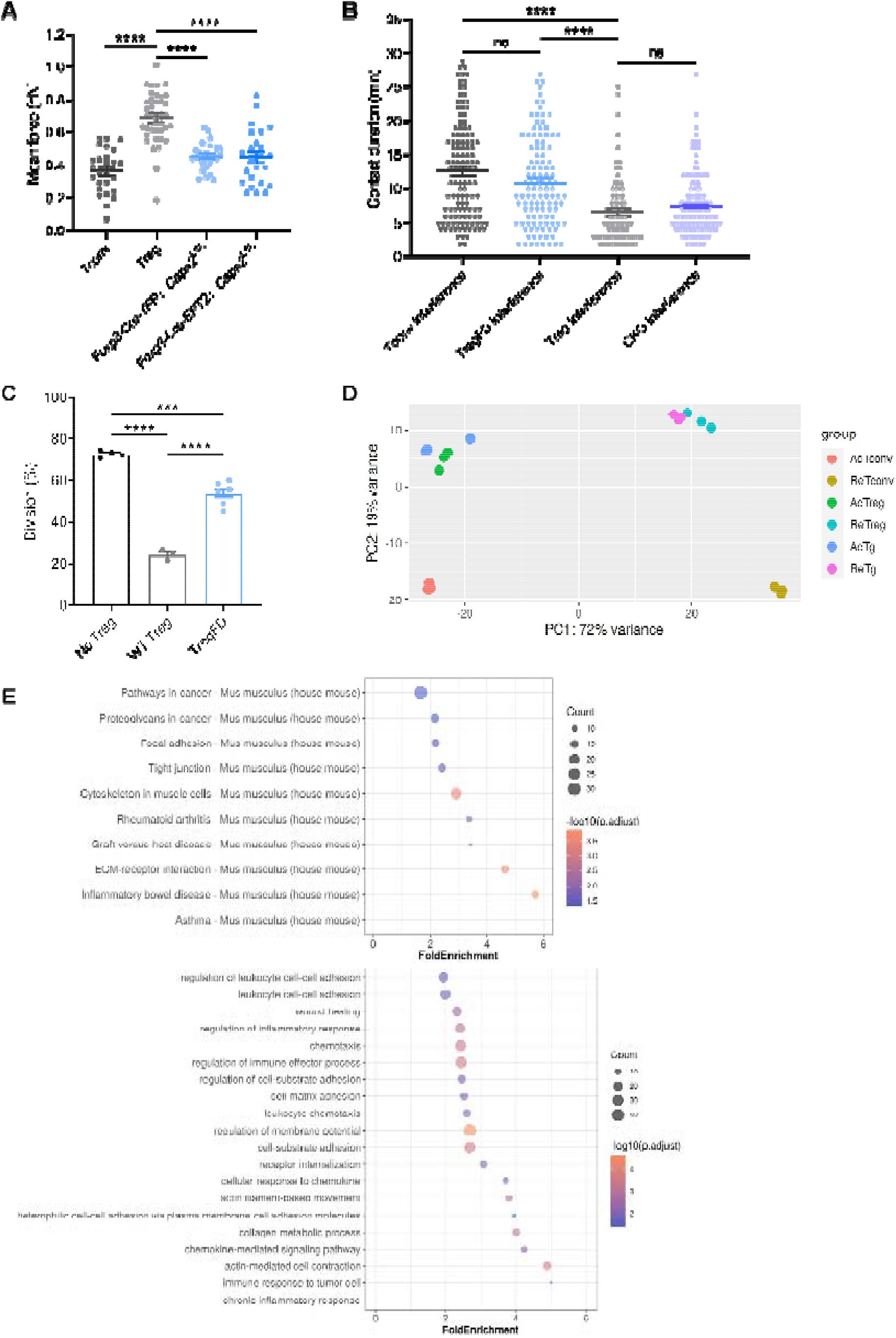
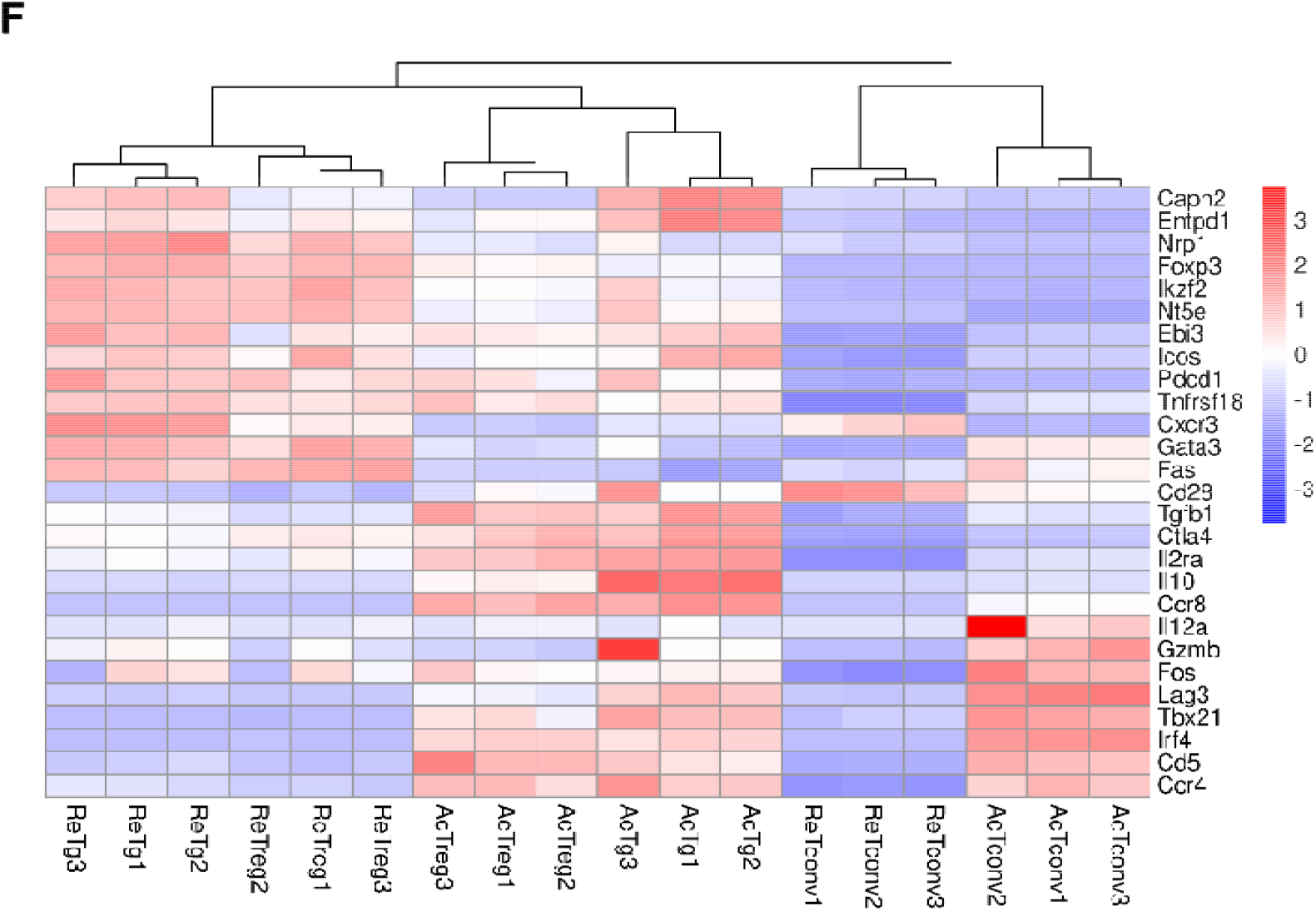
TregFD Loses DC-Mediated Antigen Presentation Blockade but Retains Treg Transcriptional Identity. (**A**) Adhesion forces between DCs and Tconv, Treg, or TregFD cells isolated from mice spleens. (**B**) In vivo imaging recorded the contact duration between DC and OT-II cells under the interference of Tconv, TregFD, Treg or RyR2 CKO cells. n = 5 mice per group; mean ± SEM; ****p < 0.0001, one-way ANOVA with Tukey’s multiple comparisons test. (**C**) In vitro suppression of OT-II proliferation. n = 3 independent experiments; mean ± SEM; ***p < 0.001, ****p < 0.0001, one-way ANOVA with Tukey’s multiple comparisons test. (**D**) PCA of bulk RNA-seq data from resting and activated Tconv, WT Treg, and TregFD cells. n = 3 biological replicates per group. (**E**) GO and KEGG pathway enrichment of DEGs between activated WT Treg and TregFD cells. (**F**) Heatmap of canonical Treg-specific suppressor genes in resting and activated Tconv, WT Treg, and TregFD cells.

### TregFD Fails to Suppress Autoimmune Diseases In Vivo

To test in vivo suppressive function, we evaluated TregFD in four autoimmune models. In DSS colitis, adoptive transfer of WT Tregs reduced weight loss, colon shortening, and histological inflammation. TregFD transfer provided no significant protection (Fig. 4A, and fig. S5A). H&E staining showed severe submucosal edema and necrosis in TregFD-treated mice, similar to model controls. In imiquimod-induced psoriasis, WT Tregs strongly reduced erythema, scaling, and skin thickening. TregFD had minimal therapeutic effect, with persistent skin pathology (Fig. 4B, and fig. S5B). In ConA-induced autoimmune hepatitis, WT Tregs prevented liver necrosis and inflammatory infiltration. TregFD failed to reduce hepatic damage (Fig. 4C). In CFA-induced rheumatoid arthritis, TregFD mice exhibited increased paw swelling, weight loss, and splenomegaly compared to WT controls (fig. S5, C and D). In all models, TregFD homed normally to inflamed tissues (fig. S5A), ruling out trafficking defects as the cause of failed suppression. Therefore, Tregs devoid of mechanical binding to DC lose immune suppression to a significant degree.

**Fig. 4.**
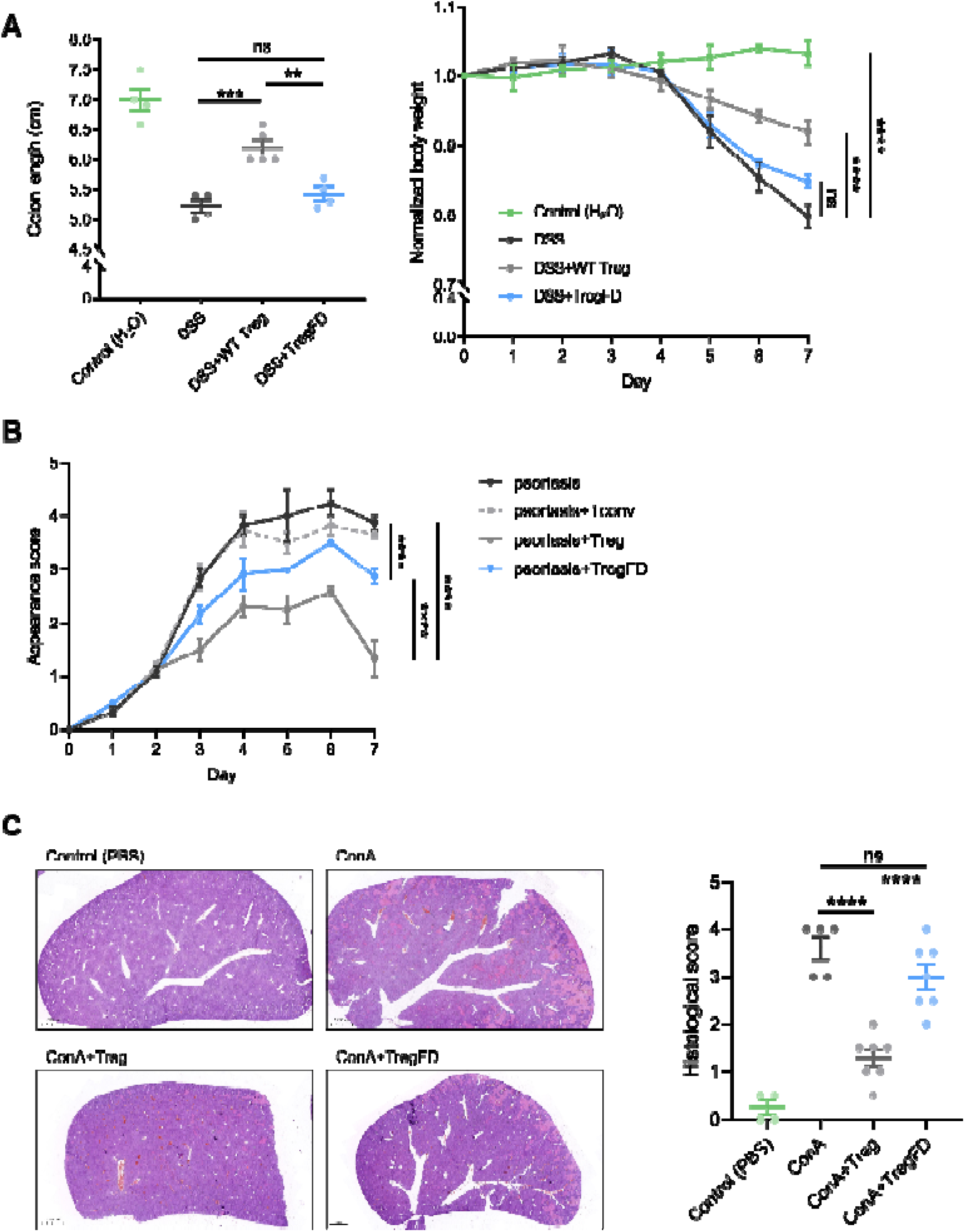
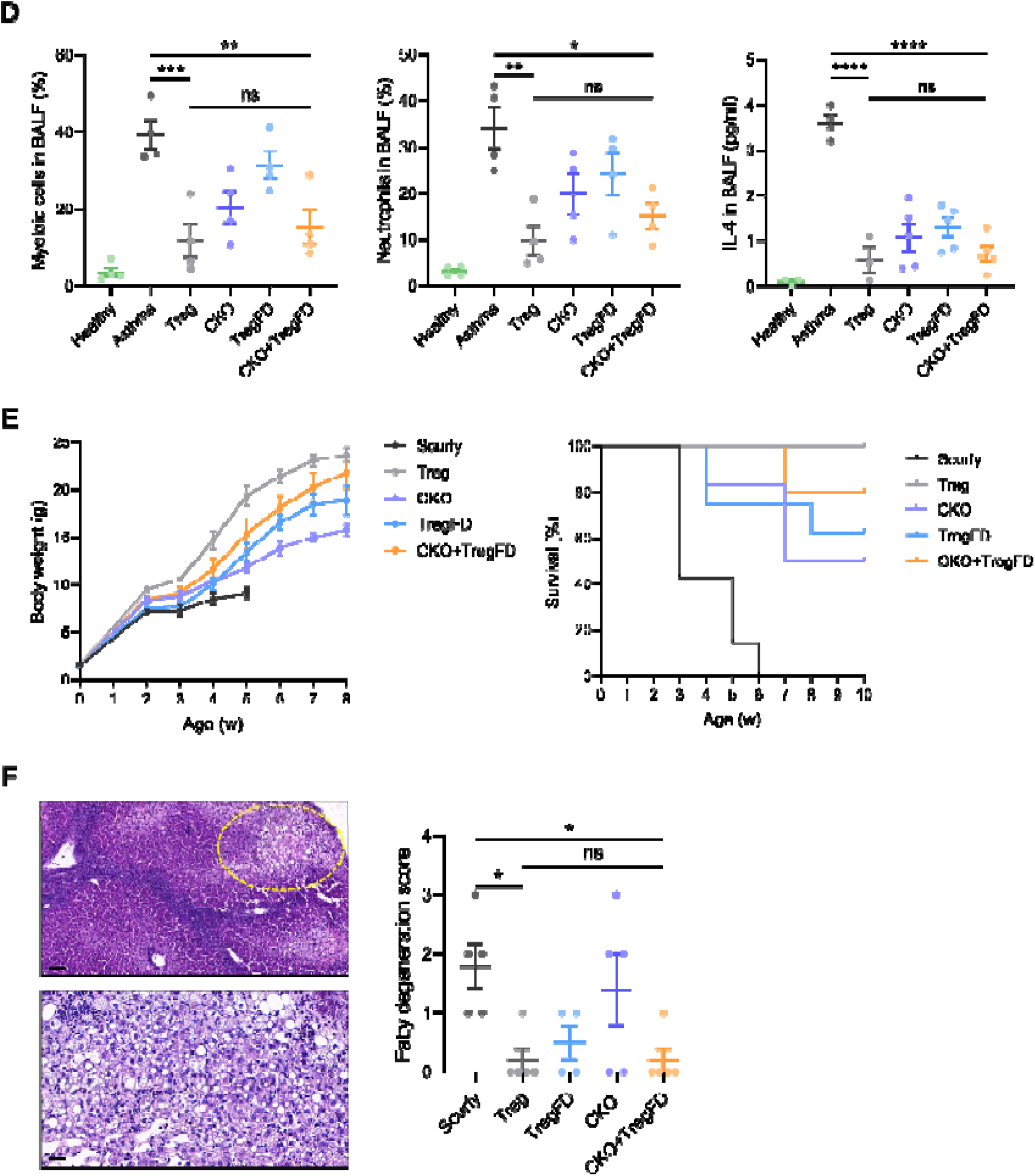
TregFD Fails to Suppress Autoimmunity; Combined Transfer with CKO Restores Function. (**A**) DSS-induced colitis: colon length (left) and percent weight loss (right). n = 6 mice per group; mean ± SEM; **p < 0.01, ***p < 0.001, one-way ANOVA with Tukey’s multiple comparisons test. (**B**) Imiquimod-induced psoriasis: clinical score over time. n = 6 mice per group; mean ± SEM; ****p < 0.0001, two-way ANOVA. (**C**) ConA-induced autoimmune hepatitis: representative liver histological sections and pathology score. n = 5 mice per group; mean ± SEM; ****p < 0.0001, one-way ANOVA with Tukey’s multiple comparisons test. (**D**) OVA-induced asthma: BALF total myeloid cells (left), neutrophils (middle) infiltration and IL-4 concentration (right). n = 6 mice per group; mean ± SEM; *p < 0.05, **p < 0.01, ***p < 0.001, ****p < 0.0001, one-way ANOVA with Tukey’s multiple comparisons test. (**E**) Scurfy mouse body weight (left) and survival curves (right) after adoptive transfer. n = 10 mice per group; log-rank (Mantel-Cox) test for survival curves; two-way ANOVA for weight. (**F**) Liver steatosis pathology score in Scurfy mice. n = 5 mice per group; mean ± SEM; *p < 0.05, one-way ANOVA with Tukey’s multiple comparisons test.

### Combined TregFD and RyR2-Deficient Tconv Cells Reconstitute Full WT Treg Suppression

RyR2-deficient Tconv cells (CKO) exhibited strong mechanical adhesion to DCs and high degree of immune protection, as reported by us previously (*17*). As TregFD lack the non-mechanical aspect of Treg immune suppression, we set to investigate how these two mechanisms are integrated. In an OVA-induced asthma model, individual transfer of TregFD or CKO partially reduced bronchoalveolar lavage fluid (BALF) immune cell infiltration and IL-4 levels. Co-transfer of TregFD + CKO reduced inflammation to the level of WT Treg transfer (Fig. 4D, and fig. S5E). In Scurfy mice, with large scale transfer of CKO Tconv cells, Scurfy mice were rescued from early lethality (*17*). However, when the cell numbers were limited (10^6^×4), differences in rescue efficiency started to appear. Neonatal adoptive transfer showed that TregFD improved survival more effectively than CKO (62% vs. 50%). Co-transfer of TregFD + CKO resulted in near-complete rescue, comparable to WT Tregs (Fig. 4E). Weight gain was also restored to WT levels in the co-transfer group. Histological analysis revealed that TregFD strongly reduced liver steatosis, while CKO preferentially suppressed skin and epithelial inflammation (Fig. 4F, and Fig. 5B). These data seem to suggest a subtle division of labor: non-mechanical suppressive modules are more involved in tissue homeostasis and repair, while mechanical modules may be more focused on blockage of antigen presentation.

**Fig. 5.**
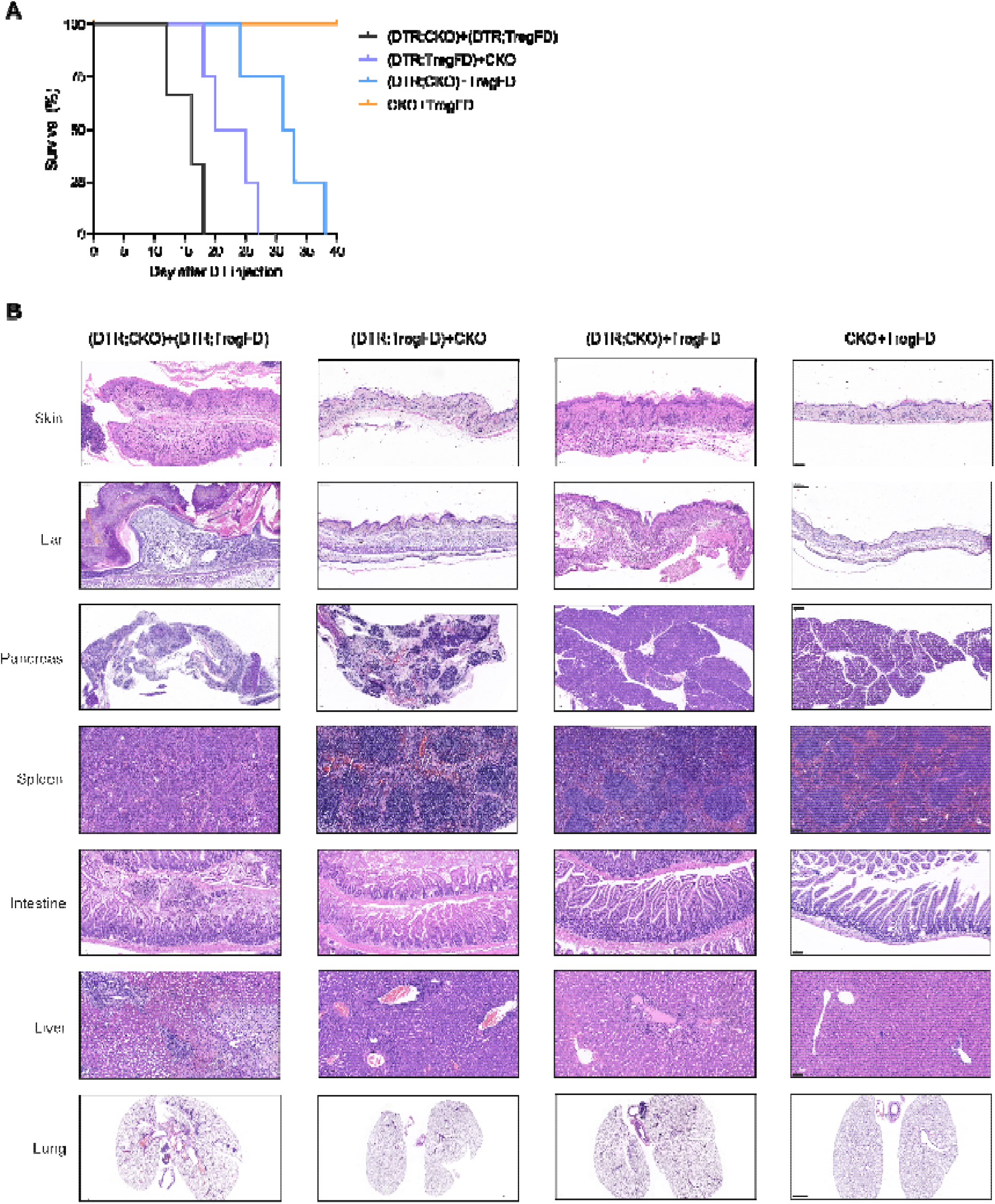
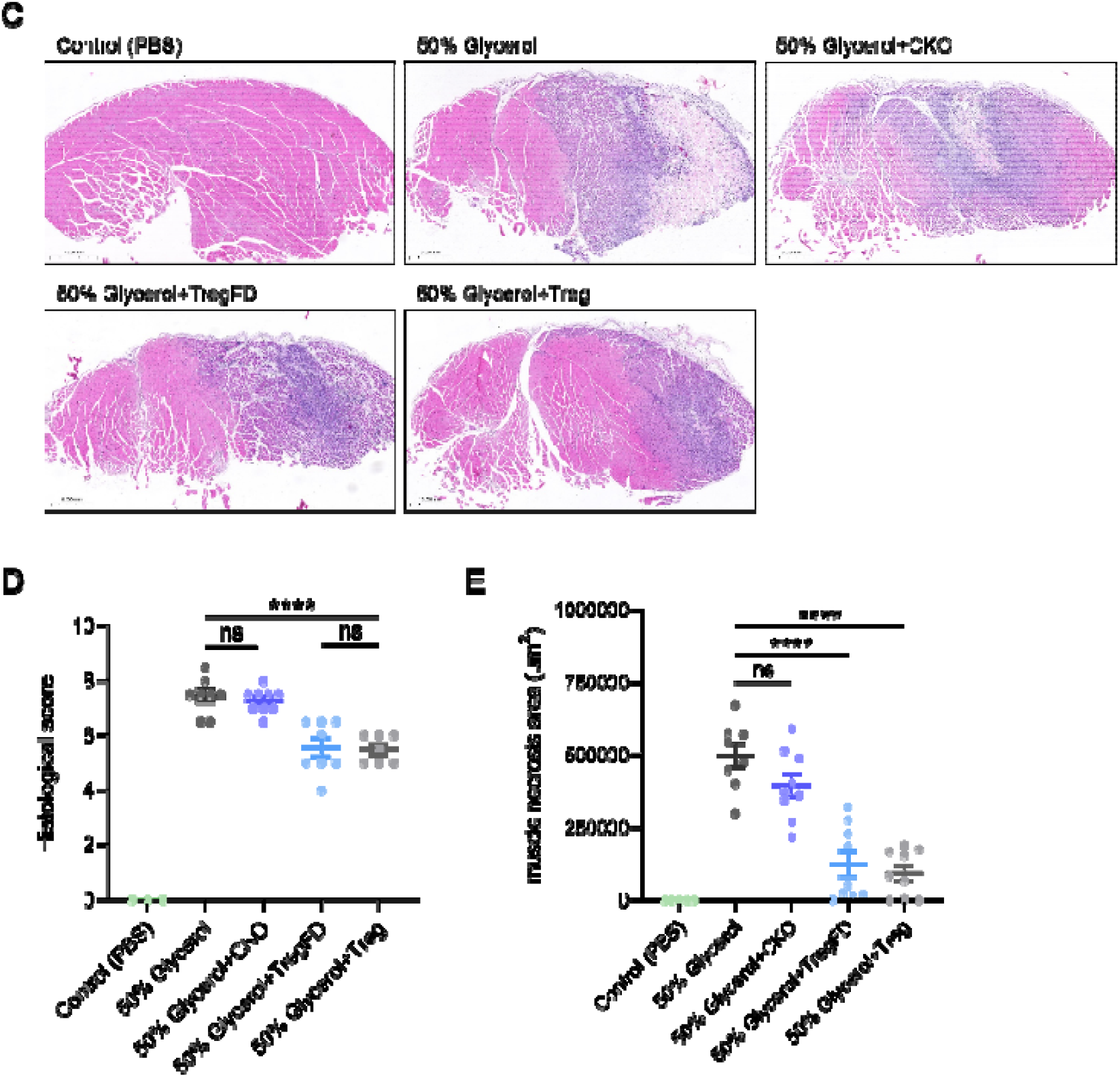
Mechanical and Non-Mechanical Treg Modules Mediate Organ-Specific Immune Homeostasis and Repair. (**A**) Survival curves of Scurfy mice after DTR-mediated depletion of CKO and/or TregFD cells. n = 8 mice per group; log-rank (Mantel-Cox) test. (**B**) Comprehensive histopathology sections of major organs. n = 5 mice per group. (**C** to **E**) Glycerol-induced muscle injury: representative H&E staining (**C**), comprehensive injury score (**D**), and muscle necrosis area (**E**). n = 5 mice per group; mean ± SEM; ****p < 0.0001, one-way ANOVA with Tukey’s multiple comparisons test.

### Mechanical Adhesion Licenses Rapid, Localized Inhibitory Cytokine Signaling to DCs

Neutralization of soluble TGF-β/IL-10 or supplementation with IL-2 did not block Treg-mediated suppression in vitro (fig. S6, A and B), indicating that these cytokines act locally rather than systemically. We hypothesized that stable mechanical adhesion promotes high-efficiency, short-range signaling. To test this, we measured activation of downstream signaling mediators in DCs. After 15 minutes of co-culture, WT Tregs induced robust nuclear translocation of pSmad2/3 (TGF-β pathway) and pStat3 (IL-10 pathway) in DCs. TregFD induced significantly weaker and delayed signaling (fig. S6, C to F). Flow cytometry quantification confirmed reduced pSmad2/3 and pStat3 frequencies in DCs co-cultured with TregFD. These results demonstrate that mechanical adhesion is a prerequisite for rapid, efficient inhibitory signaling from Tregs to DCs.

### Hierarchical Control of Immune Homeostasis and Tissue Repair

To dissect the hierarchical contributions of mechanical and non-mechanical suppression, we used DTR-mediated conditional ablation of transferred cells in Scurfy mice. This design was to create a model in which Treg suddenly lose only one of the two suppression modalities in adult stage, negating the confounding factors present in animals with immune deficiency from birth. We generated DTR;CKO and DTR;TregFD mice, allowing diphtheria toxin (DT)–mediated depletion of each module. With the full provision of both modules, mice appeared normal with no fatality. Mice lacking both modules (CKO + TregFD depleted) died earliest, within 20 days. Mice retaining only TregFD survived significantly longer than those retaining only CKO (Fig. 5A). Histopathological analysis of major organs showed that residual TregFD preserved visceral tissue integrity (pancreas, spleen, small intestine), while residual CKO suppressed inflammation in skin and ear epithelia (Fig. 5B). Mice retaining both modules showed near-healthy histology across all organs. Specifically, in ear or skin, CKO attenuated epidermal hyperplasia and inflammatory cell infiltration in the dermis. Although local epidermal hyperkeratosis was present, serous crusts were significantly fewer than in the TregFD□residual group; in pancreas, TregFD markedly alleviated acinar cell atrophy, and inflammatory infiltration was also less severe than in the CKO□residual group; in spleen, the double deletion group lacked germinal centers, indicating a marked decline in immune function. In contrast, TregFD better rescued the reduction in lymphocyte density (particularly in the red pulp) compared with CKO; as for intestine, with only CKO, the extent of mucosal necrosis was similar to that in the double depletion group, but inflammatory cell infiltration was markedly reduced. While with residual TregFD, only a small number of inflammatory cells were present in the submucosa, and small intestinal villus damage was observed in only a few mice; in lung, Scurfy mice exhibited extensive inflammatory cell infiltration around the trachea and macrophage accumulation in the alveoli. Both CKO and TregFD alleviated pulmonary inflammation, with the former showing relatively fewer inflammatory cell infiltrates.

To analyze further which module may be more involved in tissue repair (*25, 26*), we produced a glycerol-induced muscle injury model (*27-29*), TregFD reduced inflammatory infiltration, prevented fibrotic necrosis, and promoted basophilic fiber regeneration. CKO was less effective (Fig. 5, C to E). Pathological sections showed that after CKO transfer, obvious focal areas of fibroid coagulative necrosis still existed in the tibialis anterior muscle. These muscle fibers changed from tightly packed polygonal shapes to irregular rounded profiles of varying sizes, accompanied by prominent inflammatory cell infiltration in the surrounding tissue. In contrast, mice treated with TregFD exhibited smaller muscle injury areas, mainly showing regenerating basophilic fibers with central nuclei, and both inflammatory infiltration and hemorrhage were milder than those in the CKO treatment group.

To further clarify the distinct effects of mechanical and non-mechanical mechanisms mediated by Tregs on the autoimmune microenvironment in Scurfy mice, we performed single-cell RNA sequencing (scRNA-seq) on recipient CD45^+^ cells isolated from the inguinal lymph nodes of mice after DT injection. Based on the marker genes indicated in fig. S7A, endogenous immune cells were clustered into 10 distinct subpopulations (Fig. 6A and fig. S7B). The proportions of macrophages and neutrophils were significantly reduced in all treatment groups (Fig. 6B), indicating that both mechanical and non-mechanical effects contributed to the amelioration of inflammatory responses. The proportions of immune cell subsets in mice receiving Treg intervention closely resembled those observed in healthy controls. Compared with mice with residual CKO cells, littermates retaining TregFD exhibited B cell and CD8^+^ T cell proportions that were more analogous to the physiological baseline. Concomitantly, the proportion of Treg-like cells decreased to a level more comparable to that in the Treg-treated group. This phenomenon may reflect the attenuation of autoimmune inflammation, which abrogated the necessity for compensatory expansion of suppressive cell populations.

**Fig. 6.**
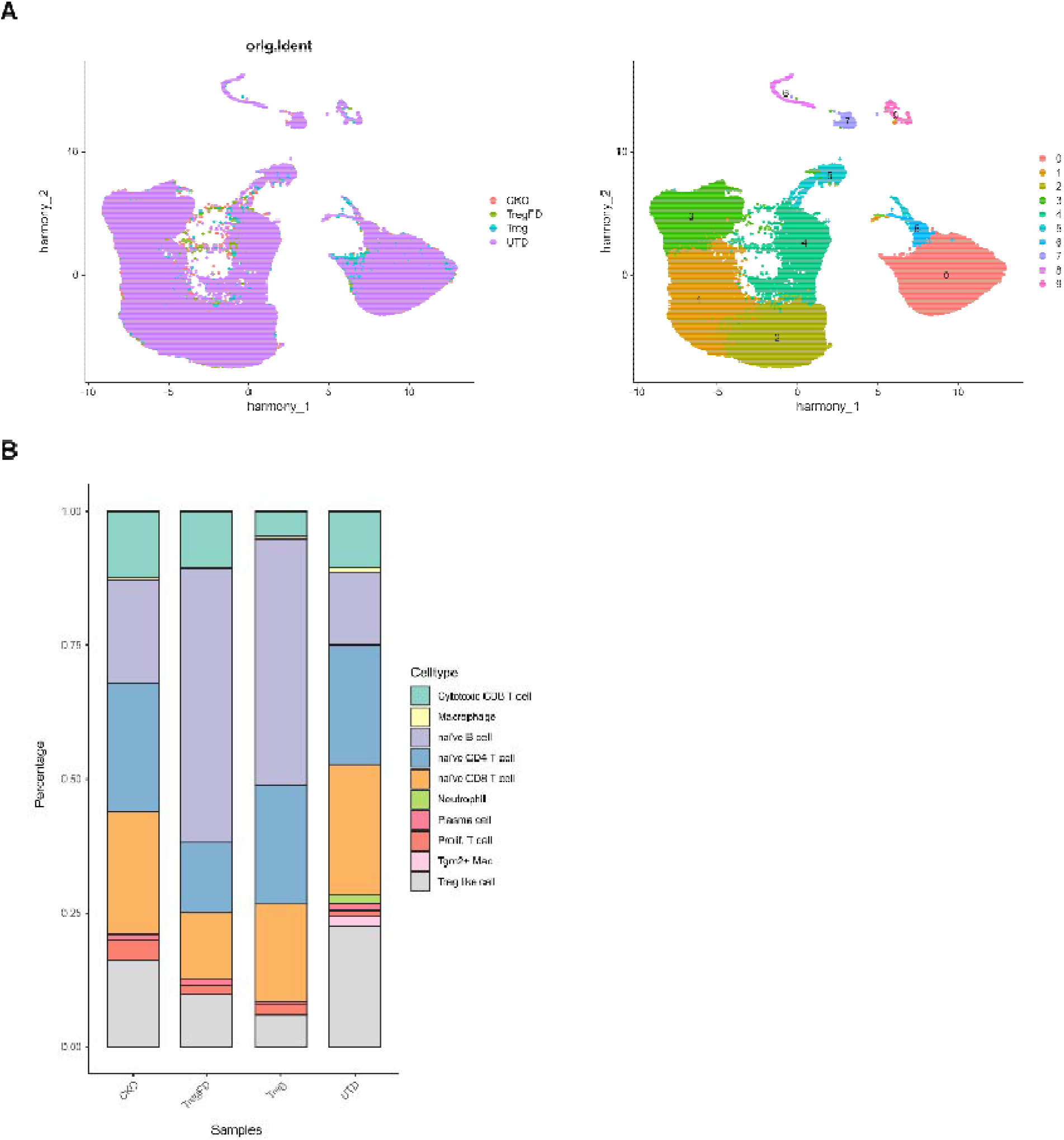
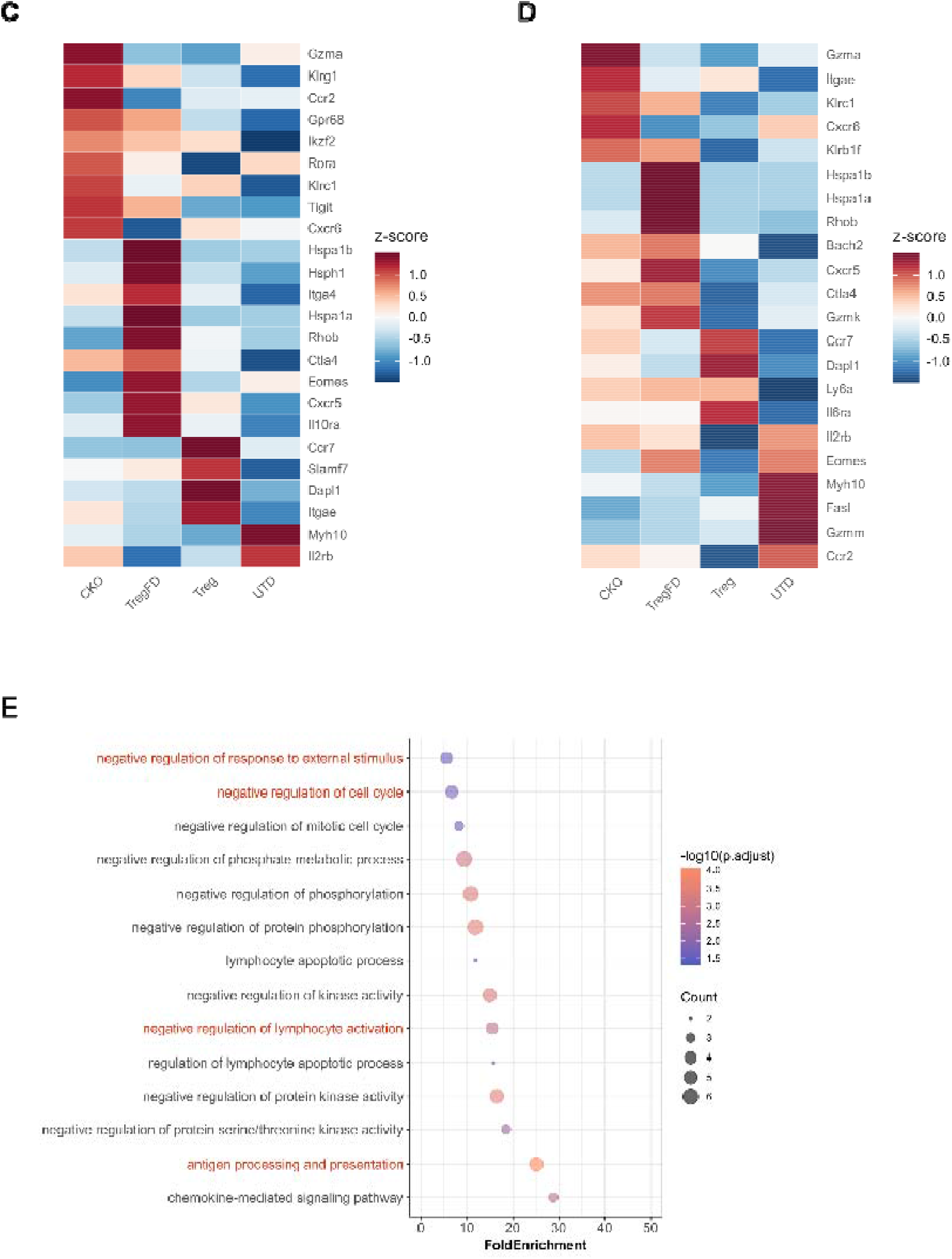
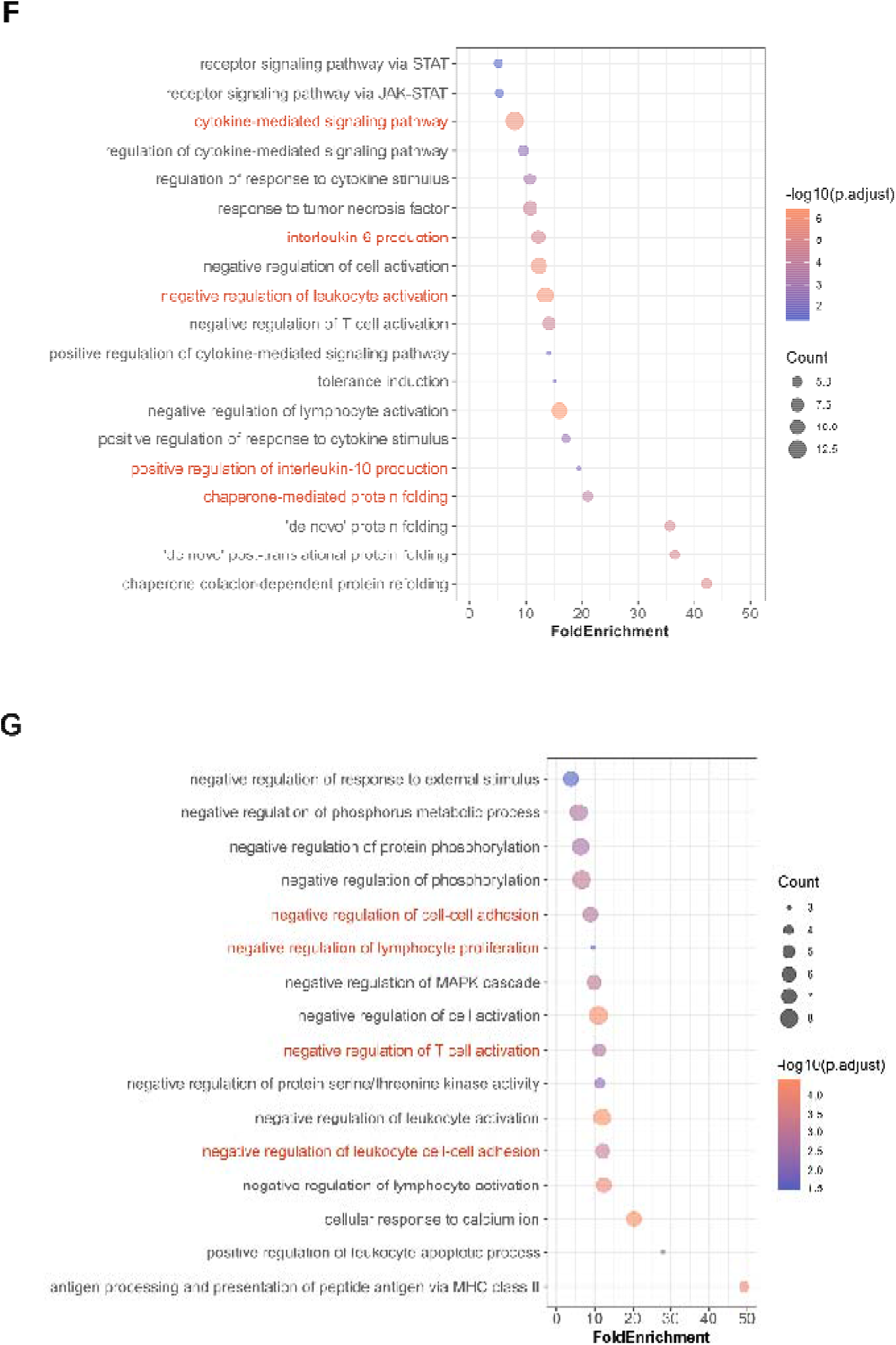
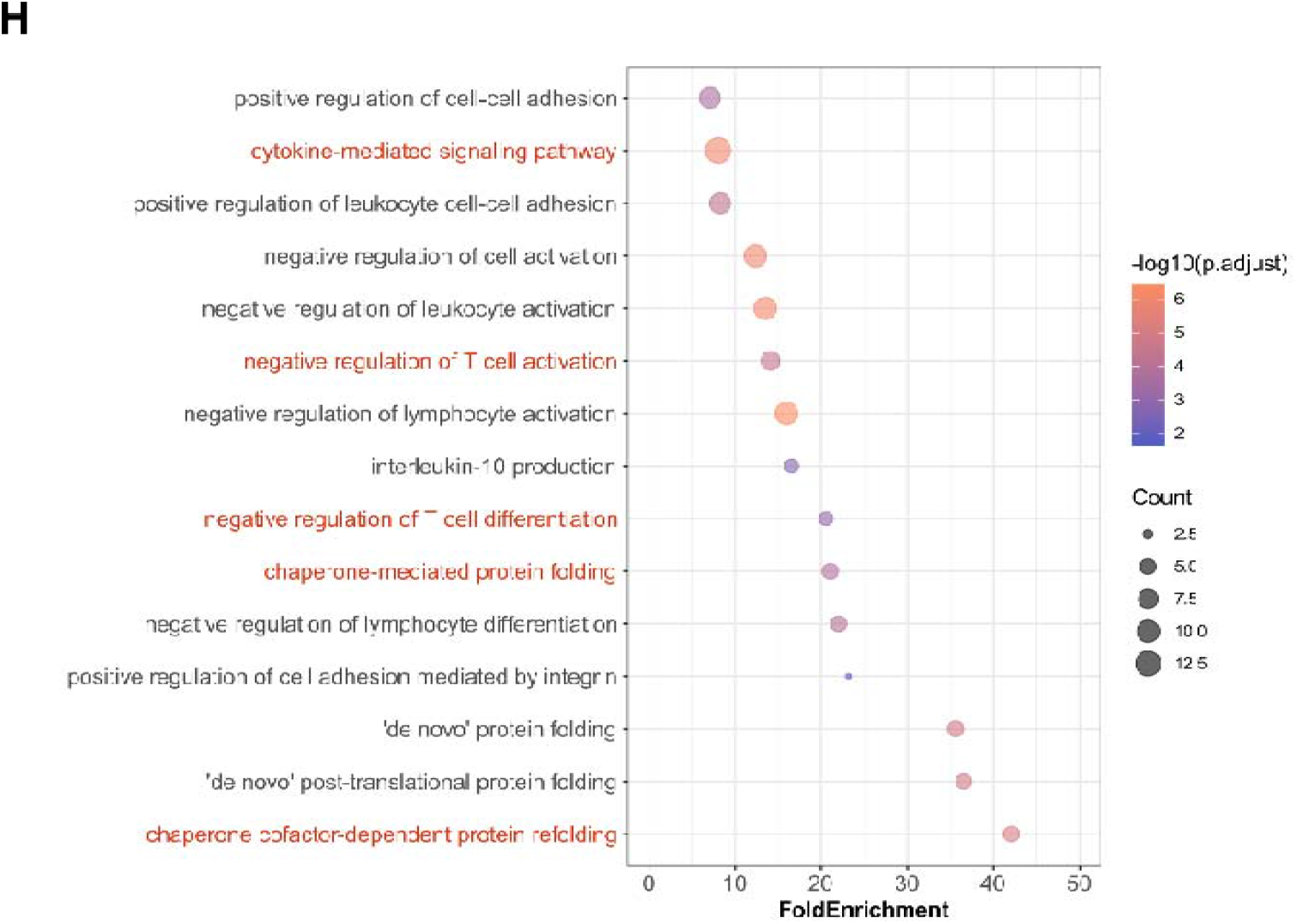
Single-cell RNA Sequencing Analysis of Endogenous CD45^+^ Cells in the Inguinal Lymph Nodes of Scurfy Mice. (**A**) UMAP visualization of CD45□immune cell subsets from Scurfy mouse lymph nodes by scRNA-seq. (**B**) Proportions of cells in each cluster in CKO-retained, TregFD-retained, Treg-retained, and untreated mice. (**C**) Heatmap of interested genes of cytotoxic CD8^+^ T cells and (**D**), naive CD4^+^ T cells under different treatments. (**E** to **H**) GO pathway enrichment of DEGs of cytotoxic CD8^+^ T cells: Treg-retaining vs TregFD-retaining (**E**), and TregFD-retaining vs untreated mice (**F**). GO pathway enrichment of DEGs of naive CD4^+^ T cells: Treg-retaining vs TregFD-retaining (**G**), and TregFD-retaining vs untreated mice (**H**).

Given that CKO, TregFD, and Treg transfer exhibited a stepwise downregulation of the proportion of cytotoxic CD8^+^ T cells in Scurfy mice, our analysis was focused on this cell population. Both mechanical and non-mechanical effects contributed to the suppression of lymphocyte activation (Fig. 6, E and F), albeit through distinct mechanisms. The former promoted CD8^+^ T cell retention in lymph nodes via adhesion-related genes such as *Itgae* and *Myh10* (Fig. 6C and fig. S7C), and also affected antigen processing and presentation (Fig. 6E). In contrast, the latter regulated the exhaustion of CD8^+^ T cells and suppressed their effector functions through cytokine-mediated signaling (Fig. 6, C and F, and fig. S7D).

CD4^+^ T cells are the major pathogenic subset in Scurfy mice (*30-32*), and they exhibited transcriptomic alteration trends similar to those of cytotoxic CD8^+^ T cells. The mechanical mechanism restricted their migration to peripheral tissues (Fig. 6D). Meanwhile, dendritic cells were sequestered by Tregs, preventing them from contacting other cells, leading to downregulation of intercellular adhesion and antigen presentation molecules in CD4^+^ T cells (Fig. 6G and fig. S7E). Conversely, the non-mechanical component primarily inhibited pro-inflammatory effector differentiation (*Bach2, Ctla4, Klrc1*, etc.) (Fig. 6, D and H) and stabilized protein folding as well as the activation state of CD4^+^ T cells via heat shock proteins (Fig. 6H and fig. S7F).

These data confirm that non-mechanical suppressive modules are dominant for systemic immune homeostasis and tissue regeneration, while mechanical modules specialize in local antigen-presentation suppression.

## Discussion

The cellular and molecular mechanisms that enable regulatory T cells to exert dominant, context-appropriate suppression have remained among the most challenging questions in immunology. In this study, we establish that mechanical adhesion as a regulatory event that controls the interaction between Tregs and DCs, and we demonstrate that this mechanical interaction is a foundational and non-redundant component of Treg suppression. By engineering a Treg-specific mechanical-deficient mouse model (TregFD), we uncouple mechanical adhesion from all other known Treg effector pathways, providing definitive causal evidence for the role of cell mechanics in immune regulation. Critically, as the mechanical binding is transcriptionally controlled by Foxp3, we illustrate a scenario whereby other non-Foxp3 tuned suppressive functions may be gathered only by Treg, rather than any other T cells to mediate a full spectrum of immune.

Our findings resolve several longstanding paradoxes in Treg biology. First, the observation that neutralization of soluble inhibitory cytokines often fails to abrogate suppression can be explained by our model: mechanical adhesion enables localized, contact-dependent cytokine delivery, rendering soluble depletion ineffective (*33-36*). Second, the in vivo dominance of Tregs over Tconvs can be attributed to their unique ability to form stable, high-force adhesive contacts that physically occupy DCs and block effector T cell priming. Third, the tissue-specific functions of Tregs can be understood through a division-of-labor model: mechanical modules control local immune synapse suppression in barrier tissues, while non-mechanical modules govern systemic homeostasis and visceral tissue repair. Mechanistically, we delineate a linear signaling cascade: Foxp3 → repression of RyR2 → reduced intracellular Ca^2^□ → inactive m-Calpain → intact Talin-1 → stabilized LFA-1 → strong Treg–DC adhesion → DC paralysis → suppression. Overexpression of constitutively active m-Calpain short-circuits this axis, making TregFD a highly specific tool for studying mechanical function. Importantly, TregFD retains all core Treg properties— development, phenotype, homeostasis, trafficking, and cytokine secretion—confirming that our model does not induce global Treg dysfunction. Transcriptomic analyses further support the purity of the TregFD model. RNA-seq shows near-identical gene expression profiles between TregFD and WT Tregs, with DEGs limited almost exclusively to cell-adhesion-related pathways. Single-cell RNA-seq in Scurfy mice extends these findings to the in vivo environment, revealing distinct immune-modulatory effects of mechanical versus non-mechanical modules. These data reinforce the idea that Treg suppression is organized hierarchically, with mechanical adhesion serving as an enabling upstream step.

Our study also has important therapeutic implications. m-Calpain inhibition may strengthen Treg mechanical adhesion for the treatment of autoimmune and inflammatory diseases. The tissue-specific division of labor we describe further allows for precision targeting: mechanical modules could be modulated in barrier tissues, while non-mechanical modules could be targeted in visceral organs. Several questions emerge for future investigation. First, how other mechanical regulators (e.g., vinculin, kindlin-3, or cytoskeletal regulators) modulate the Treg-DC adhesive complex. Second, how precisely mechanical signals integrate with metabolic and epigenetic programs in Tregs. Third, while the separation of suppression is intriguing, how Treg cells can be targeted in vivo in human settings remains a challenge. In summary, this study establishes mechanical regulation as a core pillar of Treg biology. We propose a unified hierarchical model of Treg suppression in which stable mechanical adhesion licenses and organizes downstream effector functions.

## Acknowledgments

This work was supported by the Tsinghua University Initiative Scientific Research Program and the Tsinghua-Peking Center for Life Sciences. The authors thank the Laboratory Animal Resources Center, Tsinghua University, and the Core Facility of the Institute for Immunology, School of Basic Medical Sciences, Tsinghua University, and the Imaging Core Facility, Technology Center for Protein Sciences, Tsinghua University for their services.

## Author contributions

Conceptualization: XL, YS

Methodology: XL, CH, YX

Investigation: XL, CH, XW, JK, LW

Visualization: XL, CH, CZ

Funding acquisition: YS, XW

Supervision: YS

Writing – original draft: YS, XL, CH

Writing – review & editing: YS

## Data, code, and materials availability

All transcriptional data were deposited at the NCBI Sequence Read Archive (SRA) database and are publicly available through the following BioProject IDs: PRJNA1455113 (bulk RNA-seq) and PRJNA1457244 (scRNA-seq), respectively.

