## Supplementary material for "Hierarchical Immune Suppressive Functions of Regulatory T Cells Built on Mechanical Force": Methods and Supplementary Figures

**The PDF file includes:**

Materials and Methods

Figs. S1 to S7

Materials and Methods

Mice

All mice were on the C57BL/6 background and maintained in specific pathogen‑free (SPF) conditions at the Tsinghua University Animal Center. TregFD (Foxp3‑Cre‑YFP; CAG*-*LSL‑CA‑m‑Calpain‑CFP), iTregFD (Foxp3‑Cre‑ERT2; *Capn2*^LSL^), CD11c‑GFP, CD4‑IRES‑DTR‑GFP, OT‑II, RyR2 CKO (CD4‑Cre; *RyR2*^fl/fl^; Foxp3‑GFP), Foxp3^+/−^ (Scurfy), CD45.1, and CD45.1.2 mice were bred and genotyped as previously described (Chen et al., 2017; Wang et al., 2023). All animal experiments were approved by the Institutional Animal Care and Use Committee (IACUC) of Tsinghua University.

Generation of Transgenic Mice

The CAG‑LSL‑CA‑m‑Calpain‑CFP‑WPRE construct was generated by subcloning the constitutively active mouse *Capn2* CDS (ST369/370AA) into a LSL‑containing donor vector. CRISPR/Cas9‑mediated knock‑in was performed at the H11 locus in fertilized C57BL/6 zygotes. Founder mice were screened by PCR and confirmed by Southern blot.

Cell Lines and Lentivirus Production

DC2.4 cells, L293 cells, and primary murine T cells were cultured in complete RPMI 1640 or DMEM supplemented with 10% FBS, 1% penicillin/streptomycin, and 2 mM L‑glutamine. Lentivirus was produced by co‑transfecting L293 cells with pLVX‑CMV‑IRES‑EGFP, psPAX2, and pMD2.G using polyethyleneimine (PEI). Viral supernatants were collected at 48 and 72 hours, concentrated by ultracentrifugation, and stored at −80°C.

Flow Cytometry and Cell Sorting

Spleen and lymph node cells were stained with antibodies against CD4, CD8, CD25, CD39, GITR, CTLA‑4, PD‑1, Foxp3, and appropriate isotype controls. Intracellular staining was performed using the Foxp3/Transcription Factor Staining Buffer Set (Invitrogen). CD4^+^ cells were enriched using the EasySep^TM^ Mouse CD4^+^ T Cell Isolation Kit (STEMCELL), and further sorted by flow cytometry with a BD FACSAria III or Sony MA900. Data were analyzed using FlowJo v10.

AFM‑Single Cell Force Spectroscopy (AFM‑SCFS)

Force measurements were performed using a JPK NanoWizard III AFM integrated with an inverted microscope. DC2.4 cells were plated on glass coverslips. Single T cells were attached to AFM probes coated with CellTak. Force curves were recorded at 0.5 nN setpoint and 10 s contact time. Data were analyzed using JPKSPM Data Processing software.

Intravital Imaging of Lymph Nodes

CD11c‑GFP mice were used for DC visualization. T cells were labeled with CellTrace Far Red. Cells were injected intravenously, and inguinal lymph nodes were exposed and imaged using a Dragonfly high‑speed confocal microscope. Contact duration was analyzed using Imaris software.

In Vitro Suppression Assays

DCs were sorted using the EasySep^TM^ Mouse CD11c Positive Selection Kit II (STEMCELL). OT‑II CD4⁺ T cells were labeled with CFSE. Tregs, TregFD, or CKO cells were co‑cultured with DCs and OT‑II cells at a 1:1:1 ratio in the presence of OVA_323-339_ peptide. After 60 hours, proliferation was analyzed by flow cytometry.

Bone marrow chimera

Recipient CD45.1.2 mice were administered sterile water containing 2 g/L neomycin sulfate starting from one week prior to irradiation. The two donor strains were CD45.1 Foxp3-Cre-YFP and CD45.2 Foxp3-Cre-YFP; Capn2-CFP, respectively. Recipient mice were subjected to X-ray irradiation: two doses of 6 Gy each, administered at a 3‑hour interval. Within 4 hours after irradiation, bone marrow cells were isolated from the femurs of the donor mice hind legs. The two donor cells (CD45.1 bone marrow cells and CD45.2 TregFD bone marrow cells) were mixed at an exact 1:1 ratio and intravenously injected into recipient mice. At 10 weeks post‑bone marrow transplantation, recipient mice were euthanized. The thymus, spleen, and lymph nodes were analyzed to assess the development of the two donor bone marrow cell populations into CD4⁺ T cells, CD8⁺ T cells, and Treg cells.

ELISA

Tregs, Tconvs and TregFDs were stimulated by anti-mouse CD3/CD28 antibodies for 72 hours. Cytokine levels (TGF‑β, IL‑10, IL‑2) in cell culture supernatants were measured using commercial ELISA kits according to the manufacturer’s instructions. Absorbance was read at 450 nm using a Bio‑Rad plate reader.

Western Blot

Cells were lysed in RIPA buffer containing protease and phosphatase inhibitors. Proteins were separated by SDS‑PAGE, transferred to NC membranes, blocked with 5% BSA, and probed with primary antibodies against m‑Calpain (CST), Flag (Abmart), or GAPDH (CST), followed by HRP‑conjugated secondary antibodies. Signals were detected using ECL reagent.

Bluk RNA sequencing and bioinformatic Analysis

TregFD cells were purified from Foxp3-Cre-YFP; *Capn2*^LSL^ mice, and Tconv and Treg cells were purified from Foxp3-Cre-YFP mice. After being cultured for 48 hours under either resting condition or stimulation with 5 μg/mL anti-mouse CD3/CD28, total RNA was extracted using TRIzol. Libraries were prepared and sequenced on an Illumina platform. Raw sequencing reads were quality-assessed using FastQC (v0.12.1). Adapter sequences and low-quality bases were removed using Trim Galore (v0.6.10) with the default parameters. Cleaned paired-end reads were aligned to the mouse reference genome (mm10) using HISAT2 (v2.2.1). SAM files were converted to sorted BAM format using SAMtools (v1.20) and subsequently indexed. Gene-level read counts were generated using HTSeq-count (v2.0.5) with the Gencode vM23 genomic annotation. The resulting count matrices were subsequently used for differential expression analysis.

Gene count matrices were imported into R (v4.3) and analyzed using DESeq2 (v1.42.1). Raw counts were normalized using the median of ratios method. Principal component analysis (PCA) was performed on variance-stabilized transformed data to evaluate sample clustering and identify potential outliers. Differentially expressed genes (DEGs) were identified using the Wald test with Benjamin–Hochberg correction for multiple testing. Genes with adjusted P < 0.05 and |log2 fold change| > 1 were considered significantly differentially expressed. Volcano plots were created to display the distribution of DEGs, with gene labels added for significantly regulated genes. Heatmaps were generated using normalized expression values with row-wise z-score scaling to visualize expression patterns of selected genes across experimental conditions.

Single‑Cell RNA‑Seq

Starting at postnatal day 21, mice received intraperitoneal injections of diphtheria toxin (DT, Sigma) at a dose of 0.04 μg/g body weight. Injections were administered every other day for a total of 4 consecutive doses. CD45^+^ cells were isolated from the inguinal lymph nodes of mice after DT injection. Scurfy mice with a CD45.1 genetic background were selected as recipients to exclude potential interference from CD45.2^+^ transferred cells. CD45⁺ cells from Scurfy mouse lymph nodes were sorted and processed using the BD Rhapsody single-cell multi-omics platform.

**Rawdata preprocessing, quality control and doublet detection**

Raw sequencing data were demultiplexed, aligned to the mouse reference genome (mm10, version 2020‑A, 10x Genomics) and quantified using Cell Ranger (v6.0.0) with default parameters. The resulting filtered feature‑barcode matrices were imported into R (v4.3.3) with Seurat (v5.0.3). Cells with fewer than 200 detected genes, more than 6,000 genes, or mitochondrial gene content exceeding 15% were excluded from further analysis. Percentages of mitochondrial and ribosomal protein genes were calculated using the `PercentageFeatureSet` function. Doublets were identified per sample using DoubletFinder (v2.0.4). After SCTransform normalization, principal component analysis (PCA) was performed on the top 30 PCs. Optimal pK values were selected via the BCmvn metric after parameter sweeping. Expected doublet rates were set at 8% of total cells, and homotypic doublet proportions were estimated from the clustering results. Cells classified as doublets after adjusting for homotypic doublets were removed from further analysis.

**Data integration, dimensionality reduction and cell type annotation**

Cleaned Seurat objects were merged and re‑normalized using SCTransform (v0.4.1) with percent.mt as a latent variable. PCA was performed on the scaled residuals, and the top 30 PCs were used as input for Harmony (v1.2.0) integration (`IntegrateLayers`, method = HarmonyIntegration). The harmonized low‑dimensional embedding (first 10 dimensions) was used for graph‑based clustering (Louvain algorithm, resolution = 0.15) and for uniform manifold approximation and projection (UMAP) visualisation. Cluster‑specific marker genes were identified using the `FindAllMarkers` function (Wilcoxon rank‑sum test, only positive markers, log‑fold‑change > 0.25, expressed in at least 25% of cells in either cluster, adjusted P < 0.05). Clusters were manually annotated based on canonical marker genes and visualized using DotPlot and DimPlot. The annotation was validated by cross‑referencing with the mouse cell atlas and published datasets.

**Differential expression analysis and Functional enrichment analysis**

For pairwise comparisons, the `FindMarkers` function was applied to the specified cell subsets (e.g., naïve CD8 T cell). Only genes detected in at least 10% of cells in either group and with an absolute average log₂ fold change ≥ 0.5 were tested (Wilcoxon rank‑sum test). Genes with Bonferroni‑adjusted P < 0.05 were considered differentially expressed. Volcano plots were generated using EnhancedVolcano (V1.20.0). Functional enrichment analysis was performed using the clusterProfiler (v4.19.2) package. And Gene Ontology biological processes and KEGG pathways was performed with `enrichGO` and `enrichKEGG`, respectively. Enrichment was considered significant at adjusted P < 0.05 and q < 0.2.

**Statistical analysis and visualization**

All statistical analyses were performed in R. No statistical method was used to predetermine sample size; experiments were not randomized, and investigators were not blinded to group allocation during data acquisition and analysis. Data are presented as mean ± s.e.m. or as proportions. Specific statistical tests and thresholds are indicated in the figure legends or above. Visualizations were generated using ggplot2 (v3.5.2), Seurat’s plotting functions, and custom scripts.

In Vivo Disease Models

**DSS colitis**: Mice received 4% DSS in drinking water for 7 days. Weight and stool consistency were monitored daily. Treg or TregFD cells were injected intravenously at a dose of 1×10^6^ cells per mouse on Day 1.

**Psoriasis**: 5% imiquimod cream was applied daily for 7 days. Skin erythema, scaling, and thickness were scored. Treg or TregFD cells were injected intravenously at a dose of 1×10^6^ cells per mouse on Day 2.

**Hepatitis**: One day before model establishment, Treg and TregFD cells were injected intravenously at a dose of 2×10^6^ cells per mouse. ConA (20 mg/kg, Sigma) was injected intravenously. Liver histology was scored 18 hours later.

**Arthritis**: CFA (50 μL, InvivoGen) was injected into the hind paw. Paw thickness and weight were measured daily.

**Asthma**: Mice were sensitized and challenged with OVA. On day 0 and day 14, 100 μg OVA and 4 mg aluminum adjuvant were intraperitoneally injected for sensitization. On day 21, 23 and 25, 50 μg of OVA was administered intratracheally. On day 23, Treg, TregFD or CKO cells were intravenously injected at a dose of 2×10^5^ cells per mouse. On day 32, BALF was collected for cytokine and cell analysis.

**Scurfy mouse rescue**: Newborn Scurfy mice received intraperitoneal injections of Tregs, TregFD, CKO, or mixtures (1×10^6^ cells) every 3 days for two consecutive weeks. Starting at 3 weeks after birth, diphtheria toxin (0.04 μg/g) was administered intraperitoneally. Injections were given every two days for a total of four consecutive treatments. Survival and weight were monitored.

**Muscle injury**: 50% glycerol (in PBS) was injected into the tibialis anterior muscle at a dose of 30 μL per mouse. On the following day, 5×10^5^ cells of Tregs, RyR2^⁻/⁻^ Tconvs, or TregFD were administered by intramuscular injection. Histology was performed 7 days later.

Statistical Analysis

Data are presented as mean ± SEM. Statistical tests included two‑tailed Student’s t‑test, one‑way ANOVA with Tukey’s post‑test, and log‑rank (Mantel‑Cox) test for survival curves. p < 0.05 was considered significant. Analyses were performed using GraphPad Prism 9.

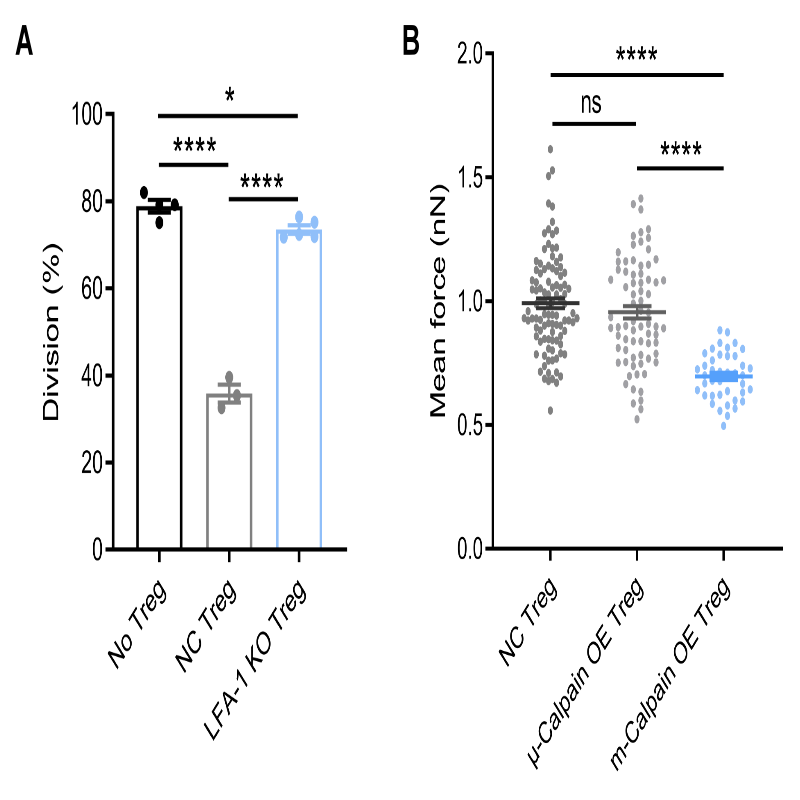

Fig. S1. Specificity of m‑Calpain in Treg Mechanical Adhesion. (**A**) In vitro suppression of OT‑II CD4^+^ T cell proliferation by WT Treg or LFA-1 knockout Treg. n = 3 independent experiments; mean ± SEM; *p < 0.05, ****p < 0.0001, one‑way ANOVA with Tukey’s multiple comparisons test. (**B**) Single‑cell adhesion force of Tregs overexpressing μ‑Calpain versus m‑Calpain. n ≥ 15 measurements per group; mean ± SEM; ns = not significant, ****p < 0.0001, one‑way ANOVA with Tukey’s multiple comparisons test.

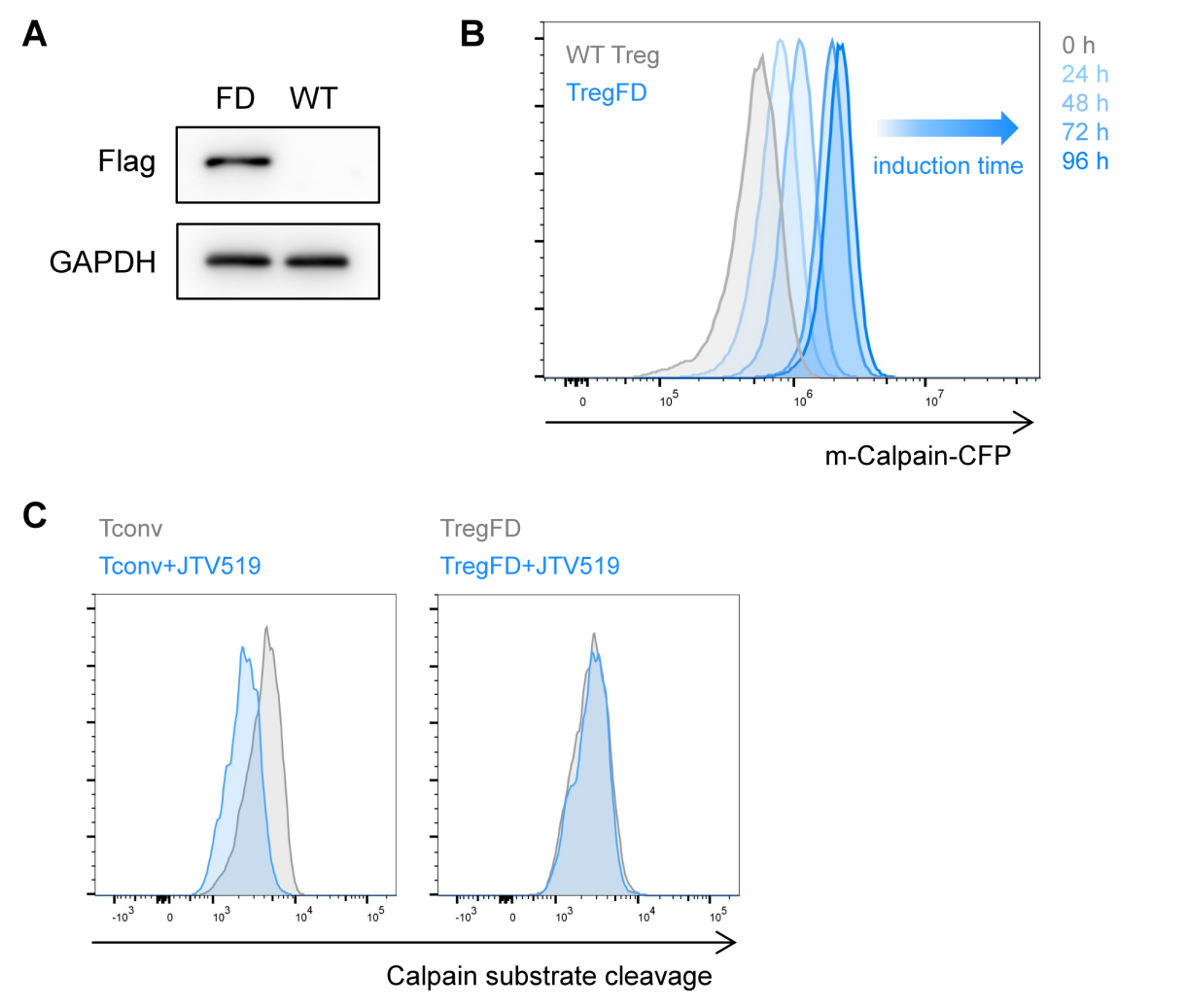

Fig. S2. Inducible Expression and Constitutive Activity of m-Calpain in TregFD. (**A**) Western blot images for Flag tag and GAPDH in WT Treg and TregFD cells. (**B**) Flow cytometry analysis of CFP induction in iTregFD Tregs at 0, 24, 48, 72, and 96 h after 4‑OHT treatment. Representative plots. n = 3. (**C**) Calpain substrate cleavage in TregFD with or without JTV-519 treatment. n = 3.

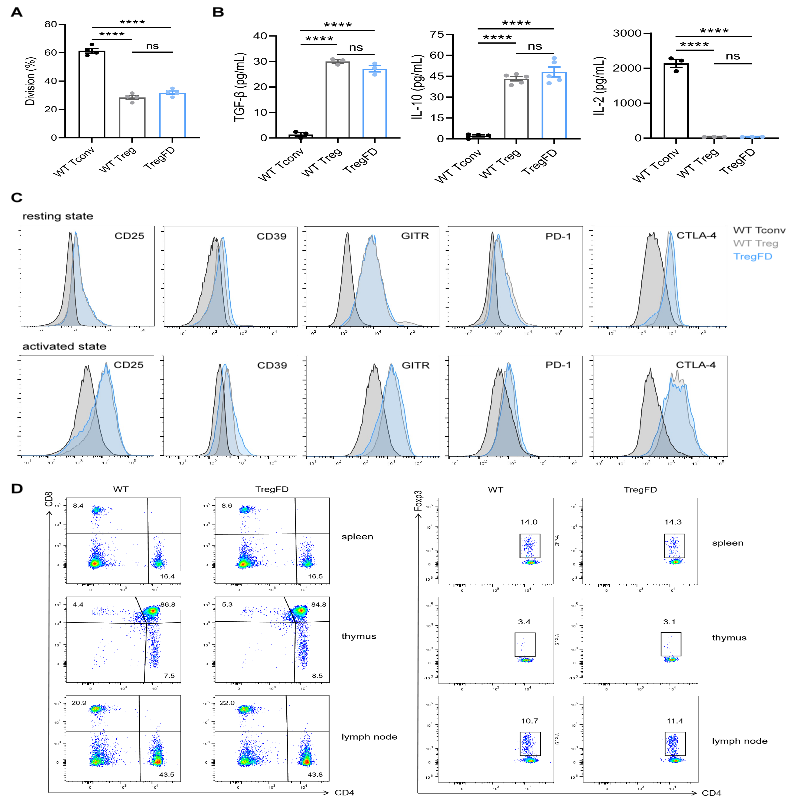

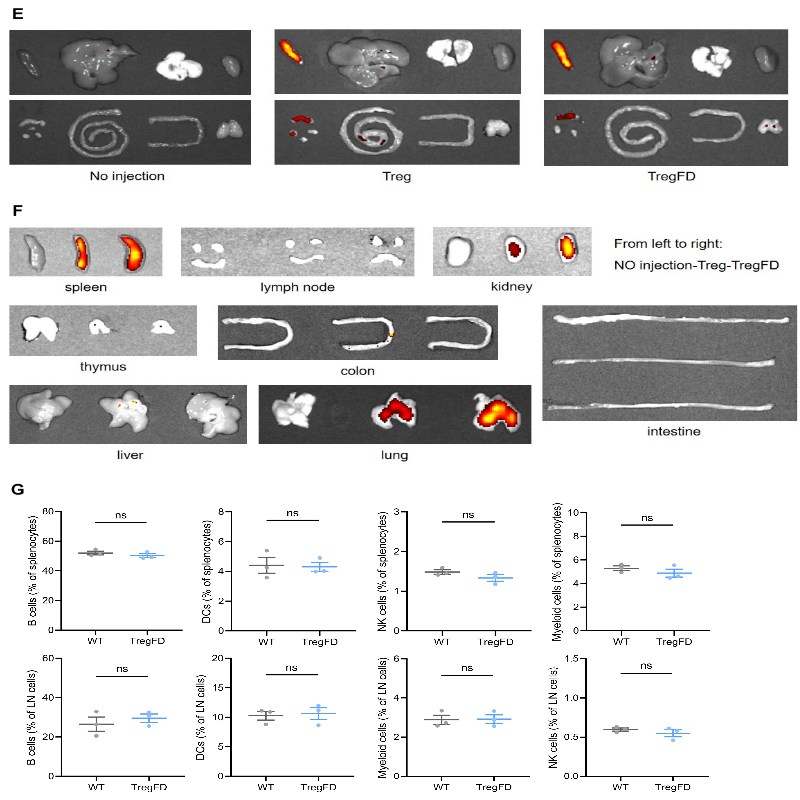

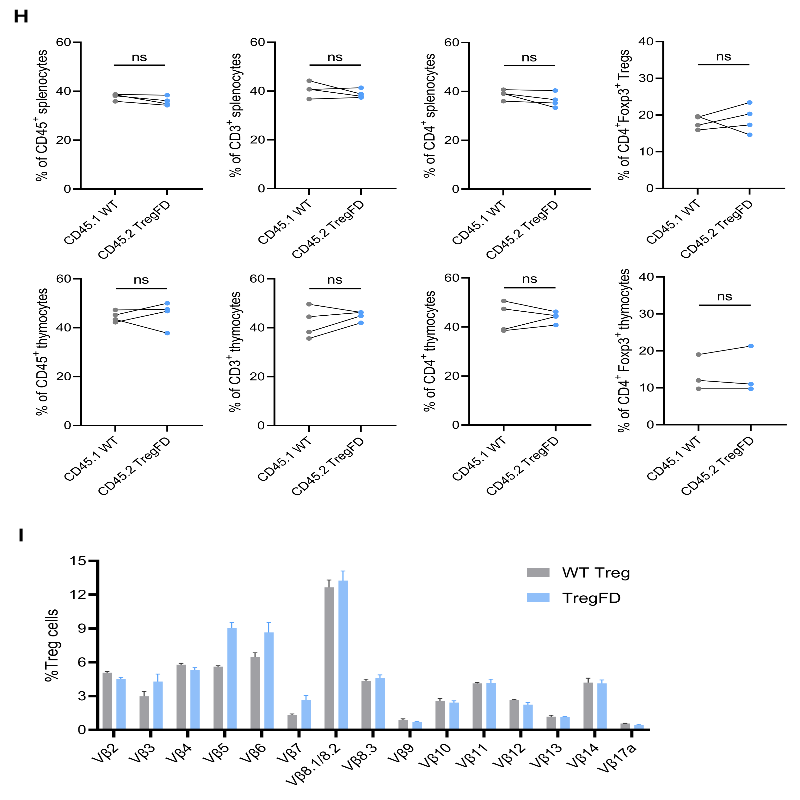

Fig. S3. TregFD Mice Are Developmentally and Phenotypically Normal. (**A**) Treg proliferation after anti‑CD3/CD28 stimulation. n = 3; mean ± SEM; ****p < 0.0001, ns = not significant, one‑way ANOVA with Tukey’s multiple comparisons test. (**B**) ELISA for TGF‑β, IL‑10, and IL‑2 in culture supernatants. n = 3; mean ± SEM; ****p < 0.0001, ns = not significant, one‑way ANOVA with Tukey’s multiple comparisons test. (**C**) Surface expression of CD25, CD39, GITR, PD‑1, and CTLA‑4 on resting and activated Tconv, WT Treg, and TregFD. Representative histograms. n = 3. (**D**) Proportions of CD4^+^, CD8^+^, and Foxp3^+^ T cells in spleen, thymus, and lymph nodes. n = 5 mice per group. (**E**) In vivo tissue trafficking of WT Treg and TregFD in healthy mice. Representative fluorescence images. n = 3 mice per group. (**F**) Trafficking in Scurfy mice. n = 3. (**G**) Proportions of B cells, DCs, NK cells, and myeloid cells. n = 5 mice per group; mean ± SEM; ns = not significant. (**H**) Bone marrow chimera reconstitution efficiency. The two points connected by the same wire come from the same mouse. n = 5; mean ± SEM; ns = not significant. (**I**) TCR Vβ repertoire diversity. n = 3.

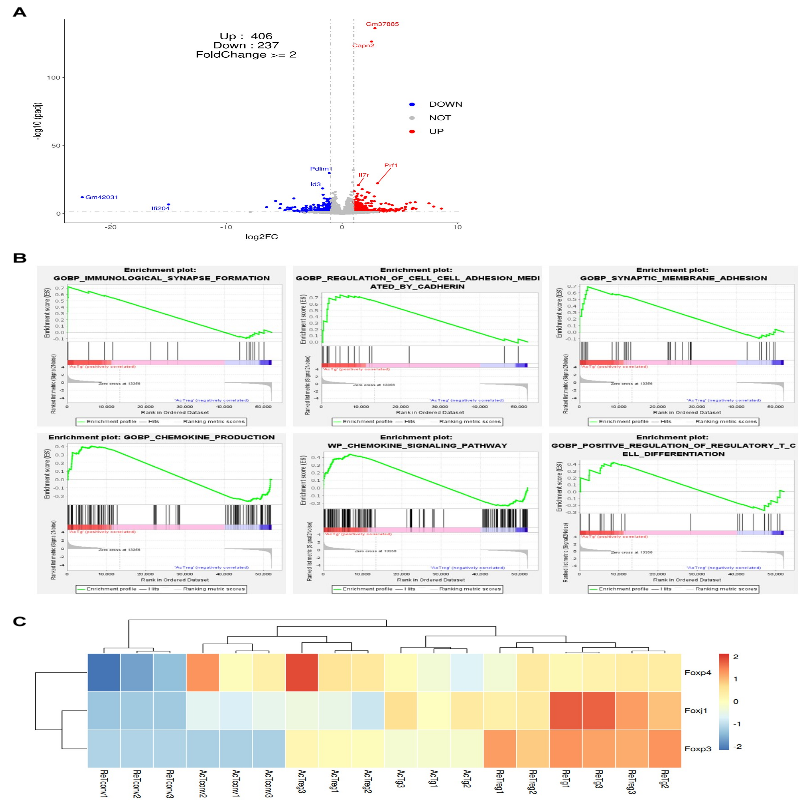

Fig. S4. **Transcriptomic Similarity between WT Treg and TregFD.** (**A**) Volcano plot of DEGs between activated WT Treg and TregFD cells. (**B**) GSEA using DEGs between activated WT Treg and TregFD cells. (**C**) Expression of FOX family transcription factors.

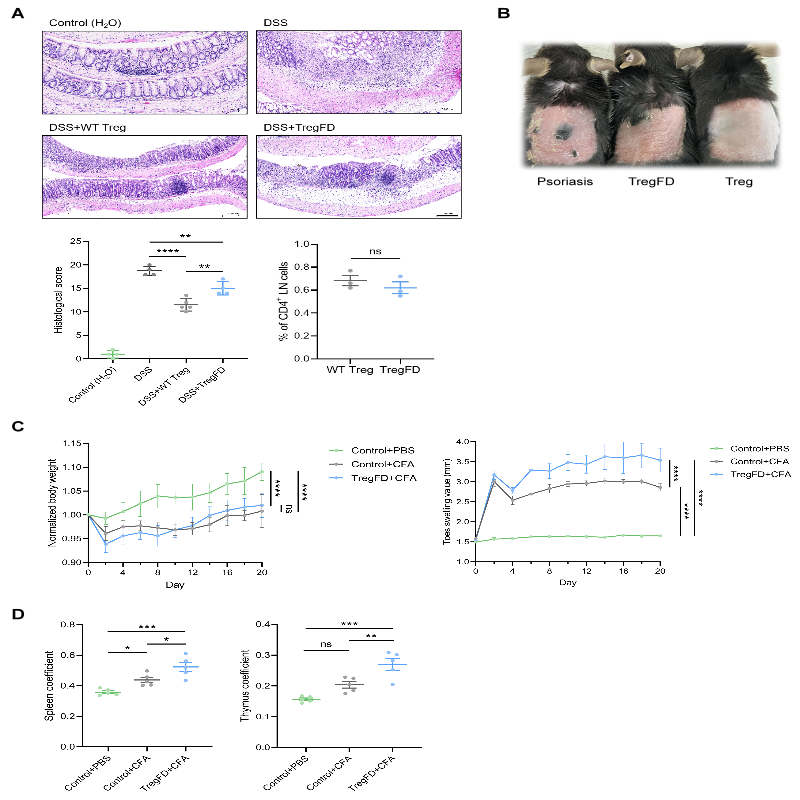

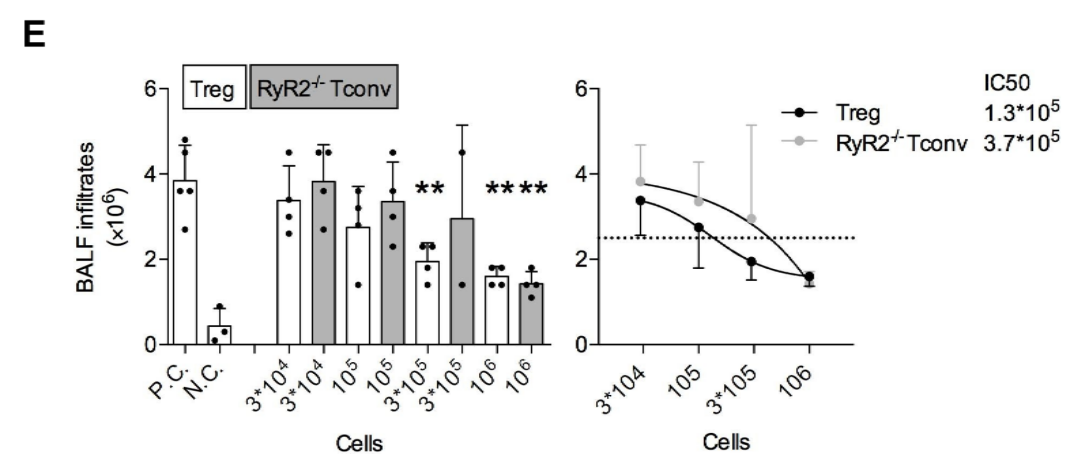

Fig. S5. **Extended In Vivo Disease Model Data.** (**A**) Representative colorectum pathological sections, histological scores, and Treg/TregFD ratios in mesenteric lymph nodes 24 hours following injection. n = 5; mean ± SEM; **p < 0.01, ****p < 0.0001, ns = not significant. (**B**) Representative skin images of psoriasis model. (**C**) CFA‑induced arthritis: weight loss and paw thickness. n = 5 mice per group; mean ± SEM; ****p < 0.0001. (**D**) Spleen and thymus coefficients. n = 5; mean ± SEM; *p < 0.05, **p < 0.01, ***p < 0.001. (**E**) Asthma model: IC_50_ comparison of WT Treg and RyR2 CKO Tconv. **p < 0.01, ****p < 0.0001.

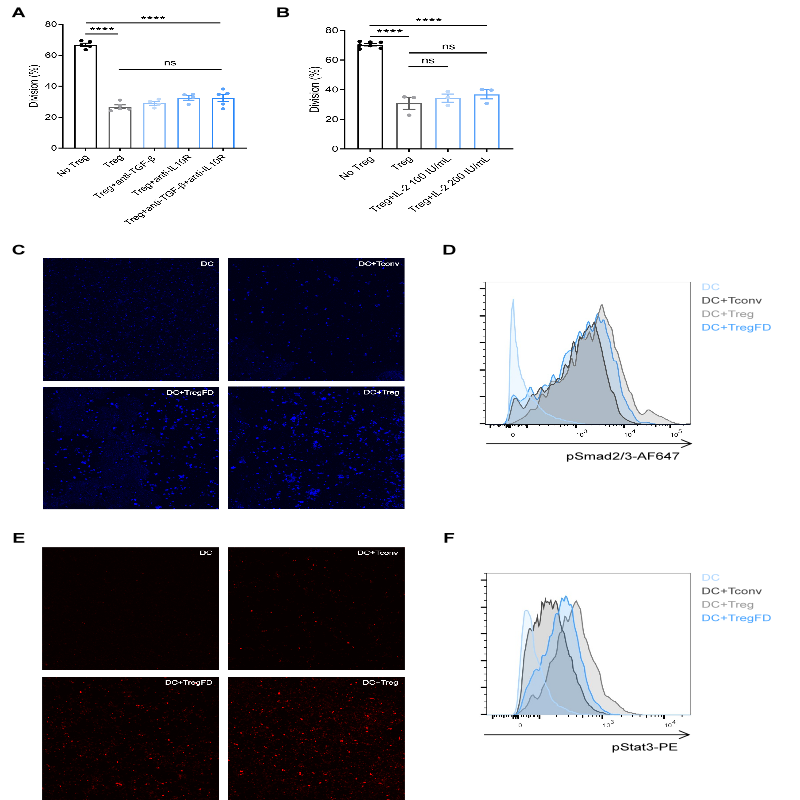

Fig. S6. **Mechanical Adhesion Licenses Rapid TGF-β and IL-10 Signaling in DCs.** (**A**) In vitro suppression with neutralizing anti‑TGF‑β and/or anti‑IL‑10 antibodies. n = 3; mean ± SEM; ****p < 0.0001, ns = not significant. (**B**) Suppression with exogenous IL‑2. n = 3; mean ± SEM; ****p < 0.0001, ns = not significant. (**C**) Immunofluorescence staining of pSmad2/3 in DCs after 15 min co‑culture. (**D**) Flow cytometry quantification of pSmad2/3⁺ DCs. n = 3. (**E**) Immunofluorescence staining of pStat3 in DCs after 10 min co‑culture. (**F**) Flow cytometry quantification of pStat3⁺ DCs. n = 3.

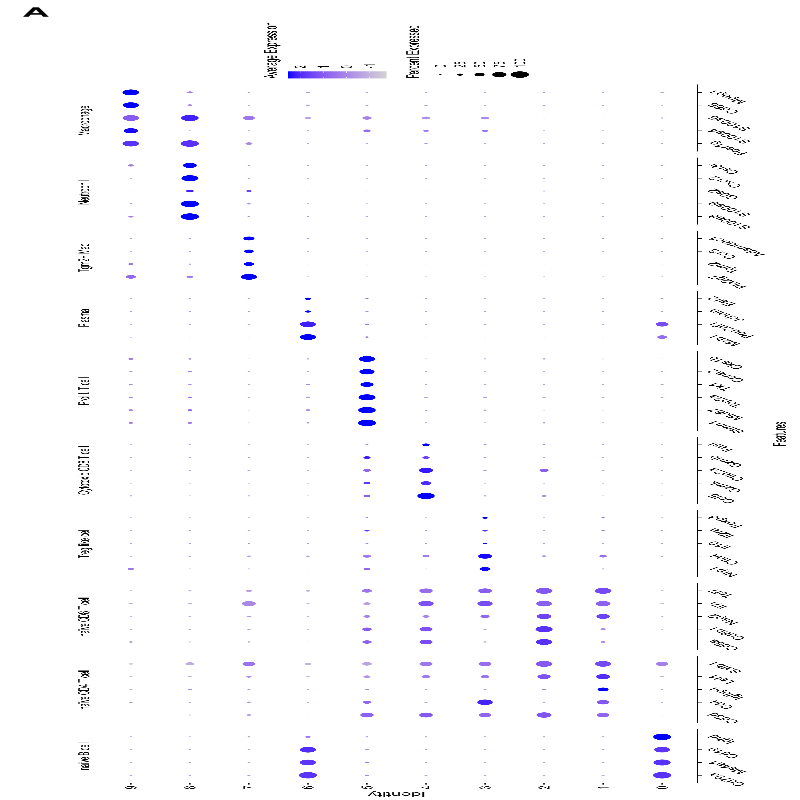

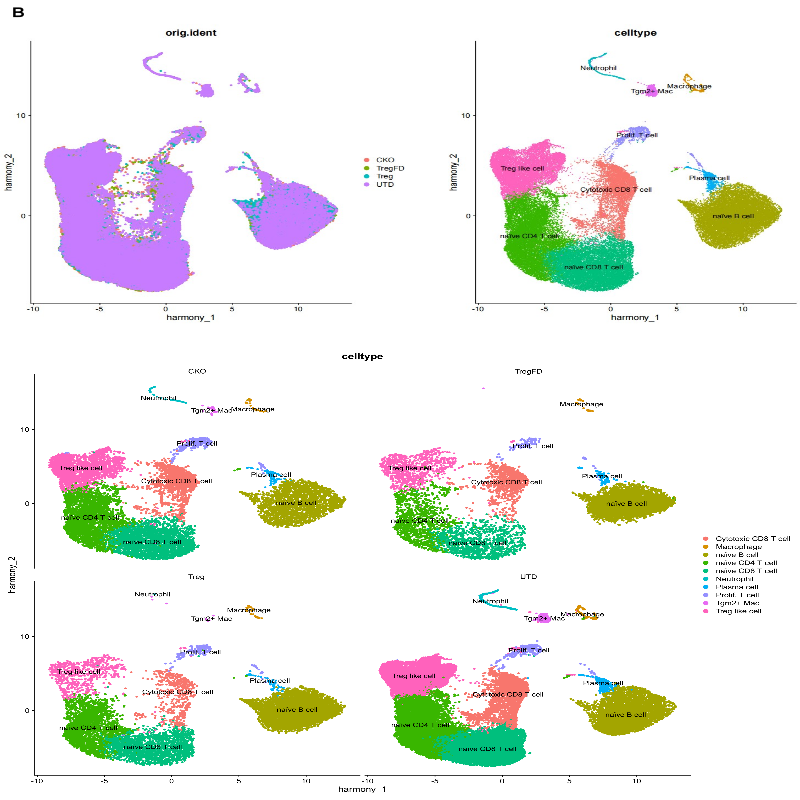

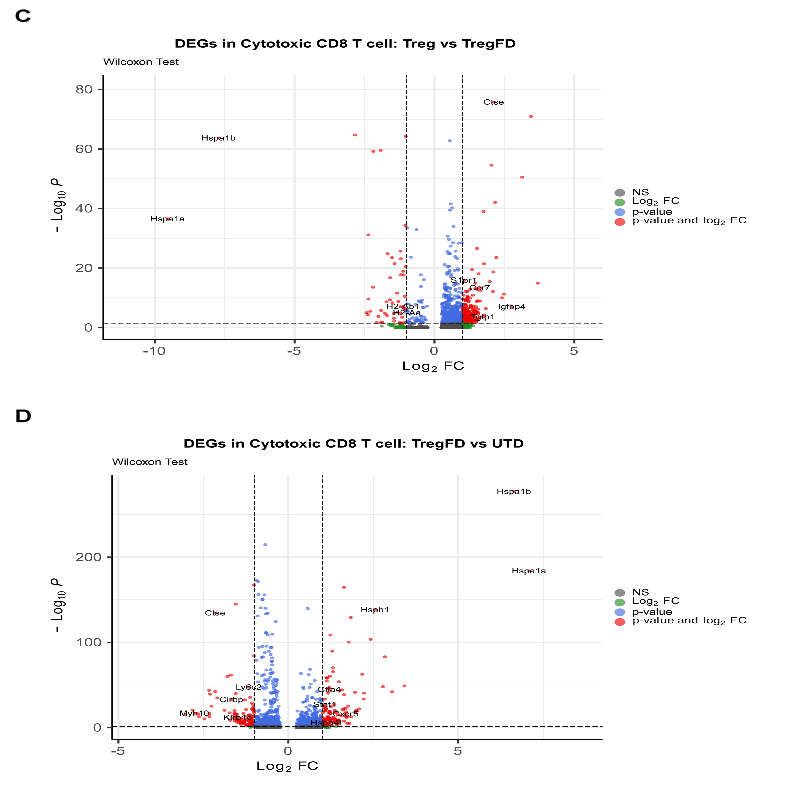

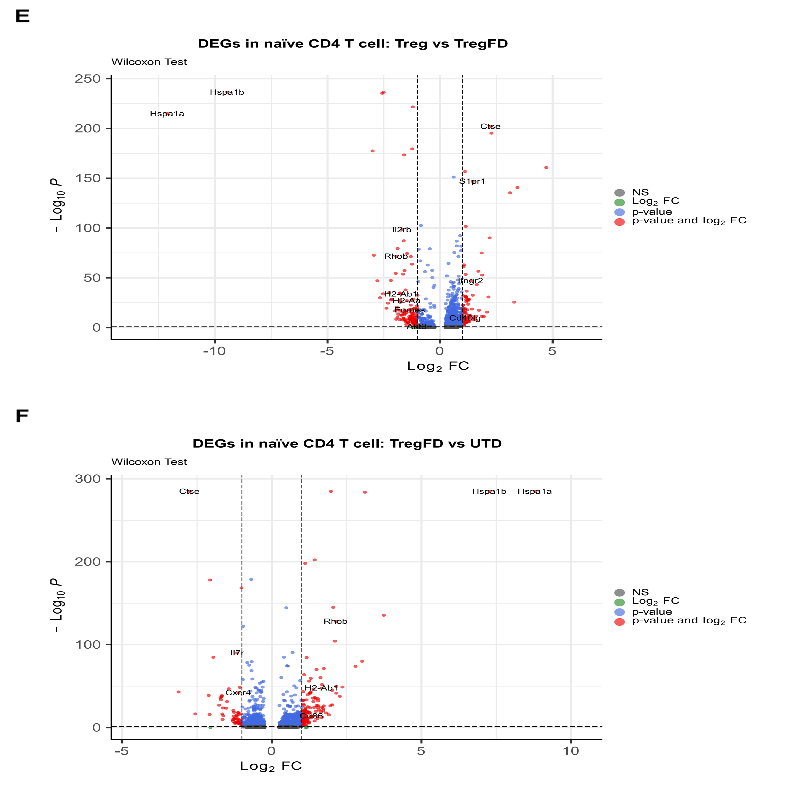

Fig. S7. **scRNA‑seq Analysis of Scurfy Mice Immune Cells.** (**A**) Canonical marker genes used for immune cell subset annotation. (**B**) UMAP with cluster labels. (**C** to **F**) Volcano plot of DEGs in cytotoxic CD8⁺ T cells: Treg‑retaining vs TregFD‑retaining mice (**C**), and TregFD‑retaining vs untreated Scurfy mice (**D**). Volcano plot of DEGs in naive CD4⁺ T cells: Treg‑retaining vs TregFD‑retaining mice (**E**), and TregFD‑retaining vs untreated Scurfy mice (**F**).
